# The *Paulinella* chromatophore transit peptide part2 adopts a structural fold similar to the γ-glutamyl-cyclotransferase fold

**DOI:** 10.1101/2025.01.15.633182

**Authors:** Victoria Klimenko, Jens Reiners, Violetta Applegate, Katharina Reimann, Astrid Hoeppner, Sander H. J. Smits, Eva C. M. Nowack

## Abstract

The chromatophores of the cercozoan amoeba *Paulinella* are photosynthetic organelles that evolved from a cyanobacterial endosymbiont. Many nucleus-encoded chromatophore-targeted proteins carry unusual N-terminal targeting signals termed crTPs. The crTPs are bipartite. Whereas crTP_part1_ that likely mediates trafficking through the secretory pathway is cleaved off during import, crTP_part2_ remains attached to its cargo protein. The function of crTP_part2_ is unknown. To contribute to unravel the functional role of crTP_part2_, here we elucidated the structures of crTP_part2_ from two different chromatophore-targeted proteins by X-ray crystallography at ∼2.3 Å resolution. Interestingly, the crTP_part2_ of both proteins adopts a structural fold. Both structures share a conserved structured core and a flexible N-terminal arm. The structured core resembles proteins of the γ-glutamyl cyclotransferase superfamily within which crTP_part2_ structures form a novel protein (sub)-family. The proposed catalytic center typical for proteins with cyclotransferase activity is not conserved in crTP_part2_. A Cys pair that is conserved in crTP_part2_ of many chromatophore-targeted proteins has been captured as disulfide bridge. Together, our data suggests that chromatophore-targeted proteins are imported in their folded state and that the fold adopted by crTP_part2_ plays a functional role during import. The characterization of its structure and flexibility provides important steps towards elucidating this novel protein translocation mechanism.

## Introduction

The transformation of bacterial endosymbionts into eukaryotic organelles has been a key process in eukaryote evolution. The only organelles identified so far that evolved by primary endosymbiosis events that were independent of the events that gave rise to mitochondria and plastids, are the photosynthetic “chromatophores” of the cercozoan amoeba *Paulinella* and the nitrogen-fixing “nitroplasts” of the haptophyte *Braarudosphaera.* In both cases, following the establishment of a cyanobacterial endosymbiont, the endosymbiont lost many functions by reductive genome evolution that were compensated by the import of nucleus-encoded proteins (Nowack and Grossman, 2012; Singer et al., 2017; Coale et al., 2024). Many of these organelle-targeted proteins carry conserved sequence extensions that apparently function as novel types of targeting signals (Singer et al., 2017; Coale et al., 2024). Their way of functioning is little understood.

In *Paulinella chromatophora*, the subject of this study, long chromatophore-targeted proteins [lCTPs; typically >250 amino acids (aa)] carry such N-terminal targeting signals that are referred to as ‘chromatophore transit peptides’ (crTPs) (Singer et al., 2017). CrTPs are ∼200 aa long, contain conserved sequence elements, and are bipartite. Upon import, crTP_part1_ is cleaved off, whereas crTP_part2_ remains attached to the N-terminus of most lCTPs (Oberleitner et al., 2022) (**Fig. 1A**). It has been proposed that the conserved hydrophobic helix in crTP_part1_ anchors crTP-carrying proteins co-translationally in the ER membrane in an N-terminus out, C-terminus in conformation and that the N-terminal adaptor protein 1 complex binding site (AP- 1 BS) is responsible for packaging lCTPs into clathrin-coated vesicles (Oberleitner et al., 2022). Although the exact timepoint at which crTP_part1_ is cleaved off is unknown, it is reasonable to assume that cleavage happens after this sorting step, possibly following fusion of the vesicles with the outer (host-derived) chromatophore membrane (Sørensen et al., 2024). This would result in a release of the cargo proteins, still attached to crTP_part2_, into the intermembrane space. The function of crTP_part2_ is unclear, but it is likely involved in mediating protein translocation across the two remaining layers (i.e., peptidoglycan (PG) and inner membrane).

**Figure 1:**
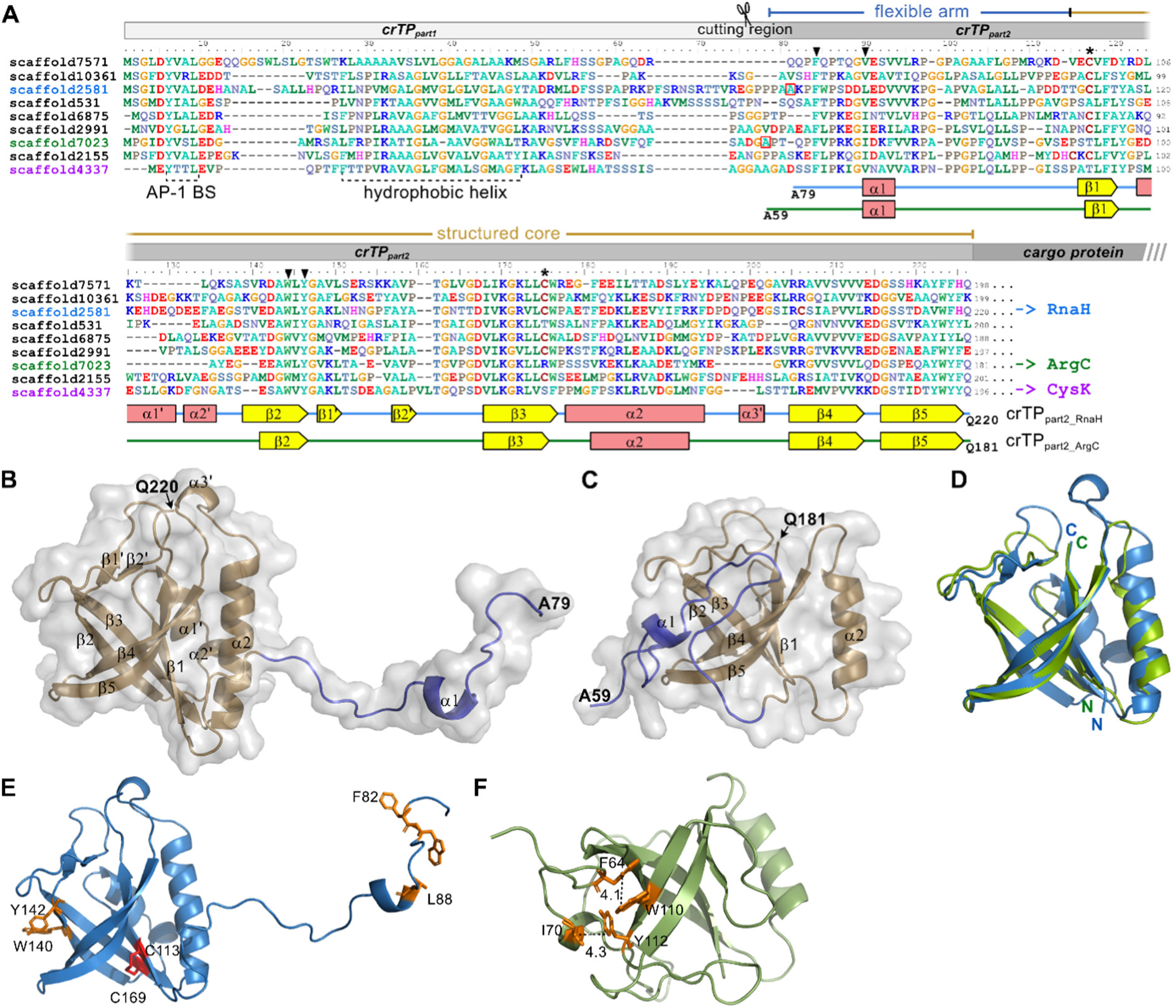
CrTPpart2 adopts a structured fold. **(A)** Multiple sequence alignment of 9 representative crTP sequences (ClustalX2, manually refined). Identifiers of proteins that were experimentally studied are highlighted in color. In crTPpart1, conserved hydrophobic helix and adapter protein complex 1 binding site (AP-1 BS) are indicated. In crTPpart2, secondary structure elements resolved by X-ray crystallography are provided underneath the alignment. Interacting Cys residues are indicated by asterisks. Conserved hydrophobic aa involved in the interaction between arm and core are marked by arrow heads. **(B-C)** Cartoon representation of crystal structure of crTPpart2_RnaH (B; pdb id 9I09) and crTPpart2_ArgC (C; pdb id 9I08). Flexible arm, blue; structured core, gold. **(D)** Superposition of the cores of crTPpart2_RnaH (blue; Leu114 to Gln220) and crTPpart2_ArgC (green; Pro93 to Gln181). **(E)** Disulfide bridge between Cys113 and Cys169 (red) stabilizes the β-barrel of crTPpart2_RnaH. In the “open conformation”, extension of the flexible arm results in exposure of hydrophobic aas (orange). **(F)** In the “closed conformation” of crTPpart2_ArgC, the flexible arm interacts with the core via hydrophobic interactions (aas marked in orange).

Interestingly, crTP_part2_ contains conserved predicted secondary structure elements across proteins (Oberleitner et al., 2022), suggesting that, different from the N-terminal transit peptides of mitochondrion and plastid-targeted proteins, which are generally unstructured, crTP_part2_ adopts a structured fold. This hypothesis guided the experimentation in this study.

## Results

### CrTP_part2_ adopts a structured fold

To contribute to the understanding of the function of crTP_part2_, we aimed to elucidate its 3D structure. For this purpose, we focused on crTP_part2_ from three chromatophore-targeted proteins. These were derived from the transcripts scaffold2581, scaffold7023, and scaffold4337 (GenBank accessions: GEZN01002575.1, GEZN01007010.1, and GEZN01004327.1, (Nowack et al., 2016)) encoding a predicted RNA helicase (RnaH), N-acetyl-gamma-glutamyl-phosphate reductase (ArgC), and cysteine synthase A (CysK); from here on crTP_part2_RnaH_, crTP_part2_ArgC_, and crTP_part2_CysK_, respectively (**Fig. 1A**). Initial analyses of these proteins by AlphaFold3 (Abramson et al., 2024) predicted the structures of the cargo proteins, RnaH, ArgC, and CysK, with high confidence; however, the crTP structures could not be modeled in high quality and resulted in largely unstructured domains (**Fig. S1**). Hence, we aimed for structure characterization by X-ray crystallography. To this end, we purified recombinant crTP_part2_RnaH_, crTP_part2_ArgC_, and crTP_part2_CysK_-containing constructs following their overexpression in *Escherichia coli* (**Fig. S2**). Two of these proteins, crTP_part2_RnaH_ and crTP_part2_ArgC_, readily formed crystals of sufficient quality to determine their structure via X-ray crystallography, at 2.4 Å and 2.2 Å resolution, respectively (for details see **Suppl. Text** and **Table S1**).

Both structures contain a structured core (colored gold in **Fig. 1B, C**) and a mostly unstructured N-terminal arm (blue in **Fig. 1B, C**). The structured cores consist of a five- stranded antiparallel β-barrel (β1-β3-β2-β4-β5) flanked by an α-helix (α2), decorated by connecting loops. In crTP_part2_RnaH_, additional very short β-strands (β1’, β2’) and α-helical elements (α1’- α3’) are embedded into the connecting loops. The N-terminal arm clearly adopts a different conformation in both structures, indicating that this part might be flexible, whereas the structured core is almost identical between crTP_part2_RnaH_ and crTP_part2_ArgC_ over a large part of the structure (rmsd 0.7 Å over 63 aligned Cα atoms, **Fig. 1D**). In crTP_part2_RnaH_, the β-barrel is stabilized by a disulfide bridge formed between Cys113 and Cys169 (**Fig. 1E**). This Cys pair is conserved in many, but not in all crTP sequences, however, if Cys occurs in these positions, it occurs as a pair (see asterisks in **Fig. 1A**). The loop connecting helix α2 with β4 is much larger in crTP_part2_RnaH_ when compared to crTP_part2_ArgC_. The loops connecting β2 and β3 as well as α2 with β3 are similar in length but adopt different conformations in the two structures, hinting towards flexibility at these positions whereas the core is rigid.

In crTP_part2_ArgC_, the N-terminal arm interacts via hydrophobic interactions between Phe64 and Trp110 as well as Ile70 and Tyr112 with the structured core and hence, shows a “closed conformation” (**Fig. 1F**). These hydrophobic residues are highly conserved between different crTP sequences (black arrowheads in **Fig. 1A**). In crTP_part2_RnaH_, for which the “open conformation” of the N-terminal arm was captured, the hydrophobic residues are exposed (**Fig. 1E**). This open conformation appears to be stabilized by the formation of a crystallographic dimer (**Fig. S3**). The N-terminal arm contains several proline residues, which give the arm a specific conformation.

To study crTP_part2_ linked to their natural cargo proteins, we purified recombinant crTP_part2_RnaH_-RnaH and crTP_part2_ArgC_-ArgC following their production in *E. coli* (**Fig. S2**). However, despite several attempts, these proteins did not form crystals.

### SAXS measurements indicate flexibility of the N-terminal arm and the linker to the cargo protein in solution

To investigate the flexibility of crTP_part2_ in solution as well as the spatial arrangement between the crTP_part2_ regions and their cargo proteins, we performed small angle X-ray scattering (SAXS) measurements. These measurements demonstrated that crTP_part2_ArgC_, crTP_part2_RnaH_, and crTP_part2_CysK_ alone are monomeric in solution (**Table S2 and Suppl. Text**). For crTP_part2_RnaH_, the “open conformation” of the flexible arm, found in the crystal dimer packing, is also present as dominant species in solution. Flexibility analyses of the termini with the Ensemble Optimization Method (EOM) recovered three potential conformations that all represent different “open conformations”, but no indication for a “closed conformation” (**Fig. 2A, B** and **Fig. S4**). In comparison, neither crTP_part2_ArgC_, nor crTP_part2_CysK_ showed flexible termini and show a more compact conformation (**Figs. S5-S6**).

**Figure 2:**
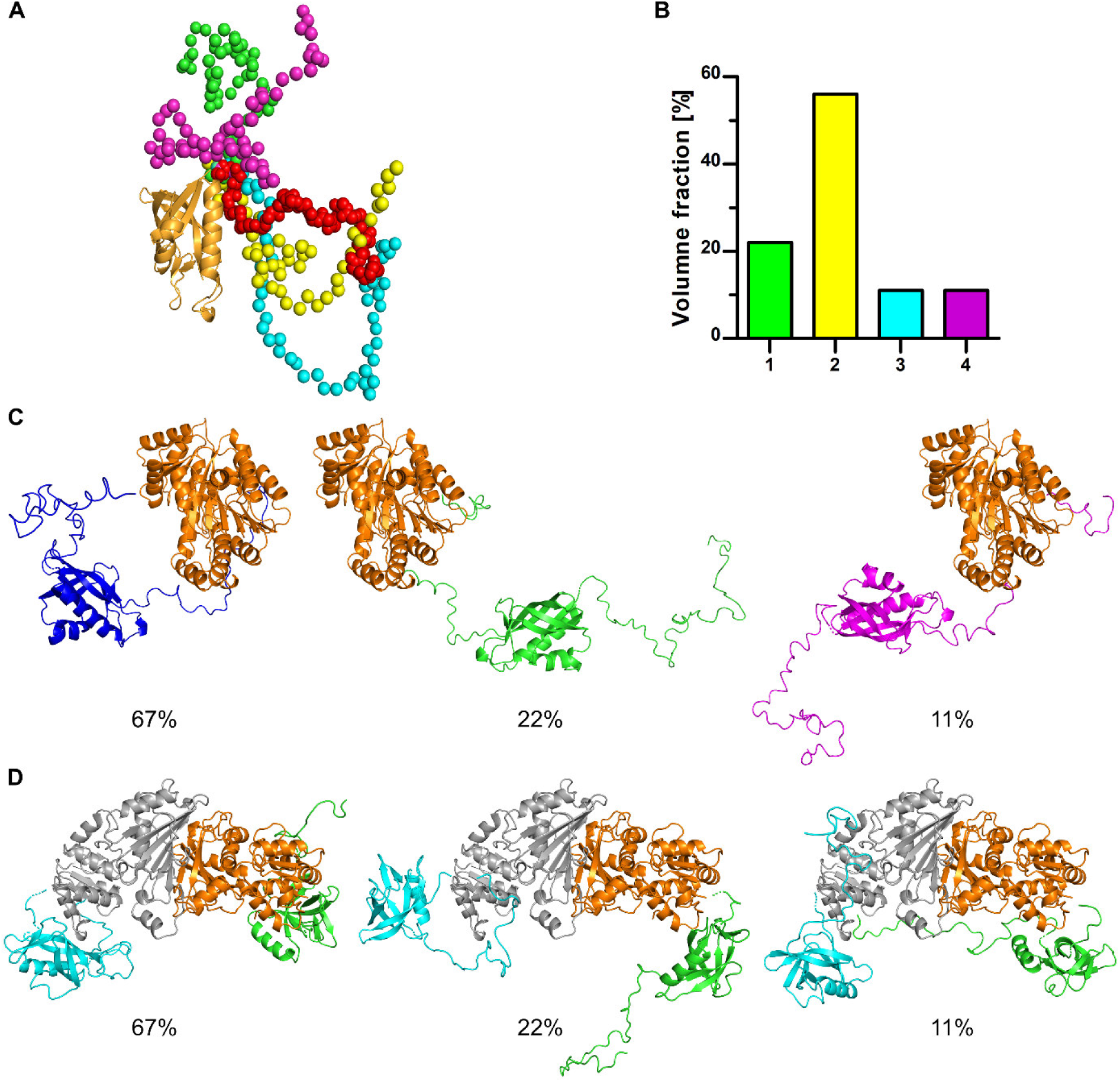
**SAXS refined models of the analyzed proteins**. **(A)** The crTPpart2_RnaH core from the crystal structure is shown as orange cartoon and the solved part of the N-terminal arm as red sphere representation. The core remained rigid for the EOM analysis and only the N and C-terminal parts were completed and used as flexible tails for the modeling. For clarity, only the N-terminal part is shown in sphere representations (in green, yellow, cyan, and magenta). **(B)** Volume fractions from the crTPpart2_RnaH EOM analysis in the corresponding color code shown as spheres in (A). **(C)** EOM models of crTPpart2_RnaH-RnaH. The RnaH core (orange) was used as rigid body. The solved crystal structure of crTPpart2 was used as flexible template and the missing linker regions were remodeled with EOM. **(D)** Dimer model of crTPpart2_ArgC-ArgC. The ArgC protomer dimer interface (grey and orange) was used as rigid body and the solved crTPpart2 as flexible template. The flexible linkers and crTPpart2 are colored in green and cyan for each protomer. The corresponding volume fractions are indicated below of each model in (C) and (D).

CrTP_part2_RnaH_-RnaH, too, is a monomer in solution (**Table S2**) and we recovered three distinct conformations, indicating flexibility between the crTP_part2_RnaH_ domain and the attached cargo protein, RnaH (**Fig. 2C** and **Fig. S7**). In contrast, crTP_part2_ArgC_-ArgC appears dimeric over the whole concentration range (**Table S2**) with the dimer interface predicted within the cargo protein ArgC. The models obtained showed that crTP_part2_ArgC_-ArgC forms an overall compact molecule, but, again, reveals flexibility between the crTP_part2_ domain and the attached cargo protein (**Fig. 2D** and **Fig. S8**). Furthermore, the N-terminal arm of crTP_part2_ArgC_ which appeared “closed” in the monomer crystal and in-solution model of crTP_part2_ArgC_ alone, now remains flexible, more in line with an “open conformation” (for more details see **Suppl. Text**).

### The structured core of crTP_part2_ shows similarity to **γ**-glutamyl cyclotransferase fold proteins

Searching the coordinates of the solved crTP_part2_ structures against the PDB database using the DALI server (http://ekhidna2.biocenter.helsinki.fi/dali/) revealed similarity of both structures to members of the γ-glutamyl cyclotransferase-like superfamily (InterPro entry IPR036568) (**Table S3**). This superfamily contains five protein families, the γ-glutamyl cyclotransferase (GGCT), the γ-glutamylamine cyclotransferase (GGACT), the glutathione-specific GGCT (GCG or ChaC), the BrtG-like, and the plant-specific GGCT-like family [**Fig. 3** and (Kumar et al., 2015)].

**Figure 3:**
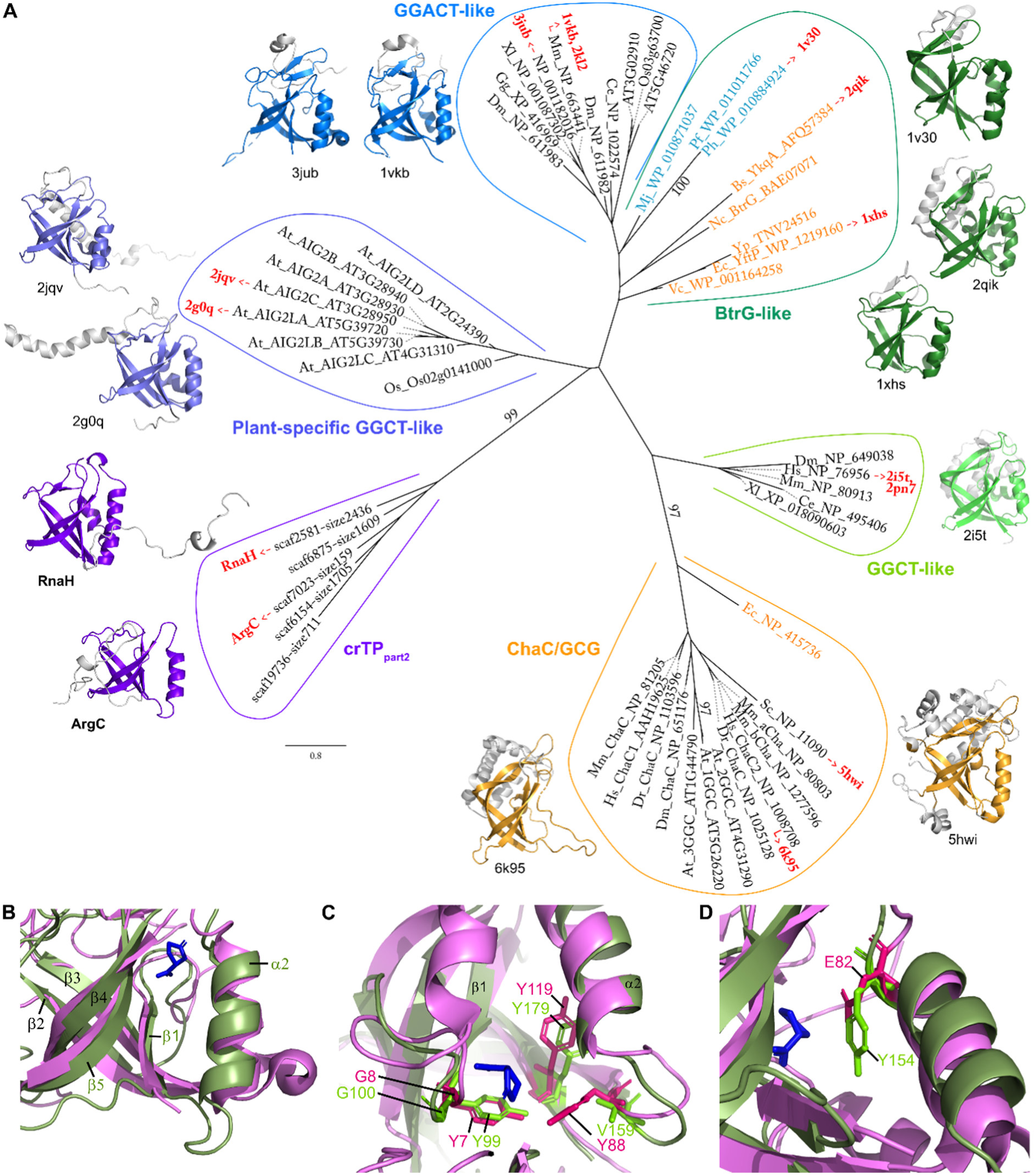
ML tree depicting the inferred phylogenetic relationship between the structured core of crTPpart2 and members of the GGCT-like superfamily. Ultrafast bootstrap values ≥95 are shown at branches. Eukaryotic sequences are in black, bacterial in orange, archaeal in blue. Protein structures available are represented as cartoon models in which the part that is structurally similar to the crTPpart2 structured core is highlighted in colors. Note that the *B. subtilis* homolog of BtrG (named YkqA, of unknown function; pdb id 2qik) contains two cyclotransferase domains within one polypeptide chain. Only the second is shown here and in Fig. S9. Species abbreviations: At, *A. thaliana*; Bs, *B. subtilis*; Ce, *C. elegans*; Dm, *Drosophila melanogaster*; Dr, *Danio rerio*; Ec, *E. coli*; Gg, *Gallus gallus*; Hs, *H. sapiens*; Mj, *Methanocaldococcus jannaschii* DSM; Mm, *Mus musculus*; Nc, *Niallia circulans*; Os, *Oryza sativa*; Pf, *Pyrococcus furiosus*; Ph, *Pyrococcus horikoshii*; Sc, *S. cerevisiae*; Vc, *Vibrio cholerae*; Xl, *Xenopus laevis*; Yp, *Yersinia pestis*. **(B-D)** Superposition of the substrate-binding pocket of the human GGACT (pdb id: 3juc, violet) in complex with the product 5-oxoproline (shown in blue) and the one of crTPpart2_ArgC (green). **(C)** Detail showing the conserved Tyr and Gly of the YGSL motif highlighted as well as other conserved or conservatively replaced hydrophobic side chains facing the substrate-binding pocket. **(D)** Detail showing the replacement of the catalytic Glu82 typical for GGCT-like proteins by a Tyr in crTPpart2_ArgC.

For 10 members of the GGCT-like superfamily structures have been experimentally solved (**Fig. 3A** and **Table S4**). Highest similarity scores were obtained for the *S. cerevisiae* glutathione-specific GGCT ChaC (pdb id: 5hwi, z-scores = 10.3 and 9.3, sequence identities = 13 and 17%), the human GGCT (pdb id: 2i5t, z-scores = 9.1 and 8.6, sequence identities = 15 and 23%), and the *B. subtilis* protein YkqA (pdb id: 2qik, z-scores = 9.2 and 8.0, sequence identities18 and 23%) (values for crTP_part2_RnaH_ and crTP_part2_ArgC_, respectively). Thus, structure comparison did not reveal affiliation of the crTP_part2_ structures to any particular family within the superfamily. In line with this result, a maximum likelihood (ML) phylogenetic analysis of a structure-guided alignment resolves crTP_part2_ sequences as a new family within IPR036568 that forms a short common branch with the plant-specific GGCT-like family (**Fig. 3A**).

For several proteins of the superfamily, enzymatic activities have been characterized and a common function of many is the cleavage of diverse γ-glutamyl derivatives by cyclotransferase activity (**Suppl. Text**). Despite their low sequence identities, catalysis is based on a similar structural fold, which includes a cavity between the β-barrel and adjacent helix that contains a conserved YGSL motif and a Glu residue that likely represents the active site (Oakley et al., 2010; Chi et al., 2014). The crTP_part2_ structures form a similar cavity build by strand β1 and β5, helix α2, and the loop between β1 and β2 that aligns with the proposed substrate-binding cavity in the human GGACT (3jub, 3juc) (Oakley et al., 2010) (**Fig. 3B**). The YGSL motif is partly conserved in crTP_part2_. Tyr99 of crTP_part2_ArgC_ at the N-terminal end of the loop connecting β1 and β2 corresponds to Tyr7 in the YGSL motif of the human GGACT. This Tyr residue is conserved throughout different crTP_part2_ sequences (**Fig. S9**) and its side chain is orientated towards the inside of the cavity (**Fig. 3C**). The following Gly is found only in around half of the crTP_part2_ sequences and is replaced in the remaining sequences mostly by other small amino acids (A, S, T, P). Ser and Leu of the YGSL motif are conserved in crTP_part2_RnaH_ but non-conservatively replaced in crTP_part2_ArgC_ by Glu and Asp. Finally, Glu82 of the human GGACT that sits at the C-terminal end of cavity-delimiting helix with its side chain oriented towards the inside of the cavity has been proposed to form the catalytic center (Oakley et al., 2010). Glu in this position is highly conserved in many proteins of the superfamily (**Fig. S9**). Interestingly, in crTP_part2_ArgC_ and crTP_part2_RnaH_ this catalytic Glu is replaced by a Tyr and an Arg, respectively (**Fig. 3D** and **Fig. S9**). In other crTP_part2_ sequences, this site harbors several other aa residues (see alignment position 192 in **Fig. 1A**). Hence, it appears unlikely that crTP_part2_ has cylotransferase activity.

## Discussion

Here we showed that crTP_part2_ domains of lCTPs in *P. chromatophora* adopt a structured fold that consists of a structured core with similarity to GGCT-like proteins and an N-terminal flexible arm that can interact via conserved hydrophobic residues with the structured core (see arrow heads in **Fig. 1A**). The proline-rich region in the N-terminal arm may represent an anchor for an interaction partner, since proline-rich regions within other proteins are known to form interaction surfaces that are responsible for interactions with e.g. elements of the cytoskeleton, peptidoglycan or biological membranes (Williamson, 1994). Interaction with a potential partner would be facilitated by the inferred flexibility of the N-terminal arm as well as the linker between crTP_part2_ and its cargo protein (**Fig. 2**). The disulfide bridge in crTP_part2_RnaH_ that has been captured by crystallography (**Fig. 1E**) is formed by a Cys pair that is conserved across crTP_part2_ domains of diverse lCTPs, which suggests that the oxidized state is biologically relevant. This assumption is in line with the oxidizing conditions in the ER lumen (Margittai et al., 2015) that has been suggested as intermediate station in the lCTP import pathway (Oberleitner et al., 2022).

Interestingly, whereas lCTPs apparently require a crTP for import, short chromatophore-targeted proteins (sCTPs; typically <90 aa) that also apparently traffic into the chromatophore via the Golgi (Nowack and Grossman, 2012) lack similar targeting signals (Singer et al., 2017). Since lCTPs and sCTPs comprise overlapping functions [e.g., cytosolic metabolic enzymes, diverse DNA-binding proteins (Singer et al., 2017; Oberleitner et al., 2020; Macorano et al., 2023)], requirement of a crTP does not seem to be tied to a specific function or final localization of the cargo protein but rather its size. This size cutoff could be set by the - so far unknown - import gate in the inner chromatophore membrane or the mesh size of the PG sacculus.

In *E. coli*, a size cutoff of ∼50 kDa has been estimated for globular proteins to be able to diffuse through the stretched PG sacculus (Demchick and Koch, 1996). Cyanobacteria generally feature a thicker PG with a much higher degree of crosslinking (Hoiczyk and Hansel, 2000). Hence, although PG composition and crosslinking has not been analyzed for chromatophores yet, it might represent an important hurdle for the transport of folded lCTPs which reach sizes >100 kDa (Singer et al., 2017). Also the plastids of Glaucophytes retained a pronounced PG layer; however, they use a TIC/TOC translocon-based mechanism for importing plastid-targeted pre-proteins in an unfolded state and only the mature stromal proteins fold into their functional conformation (Steiner and Löffelhardt, 2002). Hence, the PG does not represent a relevant size cutoff here.

Since many GGCT-family proteins cleave γ-glutamyl-containing peptides, the γ- glutamyl-containing muropeptides that cross link the PG appeared as possible ligands of crTP_part2_. Although cleavage of these peptides by crTP_part2_ appears unlikely due to the lack of conservation of the proposed catalytic center (**Fig. 3D**), we hypothesized that the binding to muropeptide derivatives could be involved in recognition of non-crosslinked areas in the PG and/or result in a conformational change enabling interaction with interaction partners at the inner membrane. However, we could not experimentally confirm binding of crTP_part2_ to the *E. coli* PG penta or tetrapeptide. Hence, if muropeptides are the natural ligands of crTP_part2_, we could not identify the exact ligand and/or correct conditions yet under which binding occurs.

In sum, our data suggests that lCTPs are imported in their folded state and that the fold adopted by crTP_part2_ plays an as of yet unknown functional role in the import process. The characterization of its structure and flexibility provides important steps towards unraveling this novel protein translocation mechanism.

## Material and Methods

### Cultivation of *P. chromatophora* and synthesis of complementary DNA (cDNA)

1. *P. chromatophora* CCAC0185 was grown as described before (Nowack et al., 2016). Total RNA was extracted and cDNA prepared as described in (Macorano et al., 2023).

### Construction of expression plasmids

The crTP_part2_ domains alone or crTP_part2_ domains plus their cargo proteins were cloned into the expression vector GPN131. This vector is a derivative of the plasmid pET-22b(+) (Novagene; 69744), in which the pelB sequence and C-terminal His_6_-tag were replaced by an N-terminal His_6_-tag, thrombin cleavage site, and SUMO-tag. For details see the **Suppl. Text**.

### Heterologous expression of recombinant proteins

For overexpression of the constructs His_6_-SUMO-TEV-crTP_part2_RnaH_, His_6_-SUMO-TEV-crTP_part2_ArgC_, His_6_-SUMO-TEV-crTP_part2_CysK_, His_6_-SUMO-TEV-crTP_part2_ArgC_-ArgC, and His_6_-SUMO-TEV-crTP_part2_RnaH_-RnaH (**see Fig. S2A**), plasmids GPN142, GPN167, GPN168, GPN195, and GPN194, respectively, were individually transformed into *E. coli* strain LOBSTR-BL21(DE3)-RIL (Kerafast, Boston, MA) (Andersen et al., 2013) and proteins were expressed under conditions detailed in the **Suppl. Text**. Finally, cells were harvested, pellets flash frozen and stored at -80°C until use.

### Protein purification

Frozen cells from expression cultures were lysed and the His_6_-SUMO-tagged proteins of interest isolated by immobilized metal ion chromatography (IMAC). The His_6_-SUMO tag was cleaved of by TEV protease and the proteins of interest purified by reverse IMAC followed by size exclusion chromatography (SEC). For details see the **Suppl. Text**. Obtained fractions were analyzed by SDS-PAGE under denaturing conditions on 12.5% polyacrylamide (ROTIPHORESE® 30; 29:1; Roth) Tris-glycine gels (Schägger, 2006) and stained with Coomassie Brilliant Blue R250 (**Fig. S2**). Protein amounts were determined by a nanophotometer (NP80, Implen).

### Protein crystallization and 3D structure determination by X-ray crystallography

CrTP_part2_RnaH_ was crystallized at 12°C with 1.5 µl of 12 mg/ml protein in buffer A (see **Suppl. Text**), mixed with 1.5 µl 23% PEG 3350 in 0.1 M HEPES pH 8.5. CrTP_part2_ArgC_ was crystallized at 12°C with 0.1 µl of 12 mg/ml protein in buffer A mixed with 0.1 µl 0.1 M HEPES pH 6.5, 2.4 M AmSO_4_ (final pH 7). Diffraction data from obtained crystals of both proteins were collected at the P13 beamline (PETRA III, DESY Hamburg) (Cianci et al., 2017). More details on experimentation, data collection, and refinement statistics are reported in **Table S1** and the **Suppl. Text**. Figures were generated using PyMOL (Schrodinger LLC; www.pymol.org).

### Small-angle X-ray scattering

SAXS data of crTP_part2_RnaH_, crTP_part2_ArgC_, and crTP_part2_CysK_ were collected on the P12 beamline at PETRA III, DESY, Hamburg) (Blanchet et al., 2015), and of crTP_part2_ArgC_-ArgC and crTP_part2_RnaH_-RnaH on our Xeuss 2.0 Q-Xoom system from Xenocs. Primary data reduction was performed with the program PRIMUS (Konarev et al., 2003). With the Guinier approximation (Guinier, 1939) implemented in PRIMUS, we determine the forward scattering *I(0)* and the radius of gyration (*R_g_*) and used the program GNOM (Svergun, 1992) to estimate the maximum particle dimension (*D_max_*) with the pair-distribution function *p(r)*. Flexible parts of the proteins were analyzed using EOM (Bernadó et al., 2007; Tria et al., 2015). Details are provided in the **Suppl. Text**.

### Phylogenetic analysis

Sequences of crTP_part2_ from indicated transcripts were aligned with diverse GGCT-like superfamily proteins downloaded from NCBI. A structure-guided alignment was generated using PROMALS3D (Pei et al., 2008). The ML tree was inferred with iqtree2 (Nguyen et al., 2015; Minh et al., 2020) using automatic model selection and 1000 ultrafast bootstrap replicates.

### Data availability

Solved protein structures were deposited in the Worldwide Protein Data Bank (https://www.rcsb.org) with accession codes provided in **Table S1**. SAXS data were uploaded to the Small Angle Scattering Biological Data Bank (SASBDB) (Kikhney et al., 2020) with accession codes provided in **Table S2**.

## Acknowledgements

We acknowledge DESY (Hamburg, Germany), a member of the Helmholtz Association HGF, for provision of experimental facilities. Parts of this research were carried out at PETRA III and we thank Tobias Gräwert and Cy M. Jeffries (EMBL Hamburg) for assistance in using beamline P12 and Gleb Bourenkov for P13.

## Author contributions

V.K., J.R., S.H.J.S., and E.C.M.N. designed the research; all authors performed research and analyzed data; V.A. and A.H. crystalized the proteins; E.C.M.N., S.H.J.S, and J.R. wrote the paper with contributions of all co-authors.

## Supplemental Results

### Structure determination via X-ray crystallography

To phase crTP_part2_RnaH_, neither the model of AlphaFold2 (Jumper et al., 2021) nor AlphaFold3 (Abramson et al., 2024) were sufficient. Therefore, we collected data from a crTP_part2_RnaH_ crystal which was soaked with potassium hexachloroiridate (IV) prior to flash-freezing, yielding a 2.6 Å dataset which was used for initial phasing and model building. Once the backbone was built, we used this model to phase the high-resolution dataset of crTP_part2_RnaH_ using molecular replacement. This model was refined to 2.4 Å. The final structure of crTP_part2_RnaH_ served as search model to obtain phases for the 2.2 Å crTP_part2_ArgC_ (data statistics are shown in **Table S1**).

### SAXS measurements indicate monomeric form of crTP_part2_ in solution and flexibility of its N-terminal arm and linker to the cargo protein

SAXS measurements demonstrated that crTP_part2_ArgC_, crTP_part2_RnaH_, and crTP_part2_CysK_ (the latter did not form a crystal) are monomeric in solution (**Table S2**). The crTP_part2_RnaH_ data revealed an elongated particle in the *p(r)* function and flexibility of the termini in the dimensionless Kratky plot (**Fig. S4**), which indicates that the “open conformation” of the flexible arm, found in the crystal dimer packing, is also present as dominant species in the monomeric in-solution form. Based on the solved core part of crTP_part2_RnaH_ we analyzed the flexibility of the termini with the Ensemble Optimization Method (EOM) (χ^2^ of 1.062) to cover potential conformations (**Fig. 2A**). All models show an “open conformation” with different conformations, but no indication for a “closed conformation”. In comparison, neither crTP_part2_ArgC_, nor crTP_part2_CysK_ showed flexible termini in the dimensionless Kratky plot (**Figs. S5-S6**) and indicate a more compact conformation. Some terminal residues were missing in the crystal structure of crTP_part2_ArgC_ so we added these residues with CORAL (Petoukhov et al., 2012) to the solved crystal structure, however, only when we allow flexibility of the whole tail (residues 1-40), we could archive a good agreement (χ^2^ of 1.293) with the experimental data (**Table S2**). The result is a slightly different position of the tail, but still in the “closed conformation”. Since crTP_part2_CysK_ did no crystallize and neither AlphaFold2 (Jumper et al., 2021) nor AlphaFold3 (Abramson et al., 2024) were able to predict a model with a high plDDT score (Mariani et al., 2013), we calculated a homology model with SWISS-MODEL (Waterhouse et al., 2018) based on the solved crystal structure of crTP_part2_ArgC_ and remodeled the N-terminal arm (**Fig. S6**, **Table S2**). Nevertheless, the best model obtained reached only a χ^2^ of 1.768 and the corresponding error plot showed mismatches over the whole range of the data (**Fig S6, Table S2**), precluding a clear interpretation of the data.

To understand the spatial arrangement and flexibility between the crTP_part2_ and its ‘cargo protein’, next, we performed SAXS measurements on crTP_part2_RnaH_-RnaH and crTP_part2_ArgC_-ArgC. The resulting data revealed that crTP_part2_RnaH_-RnaH is a monomer in solution over the concentration range measured (max. 9 mg/ml) (**Table S2**). We generated a hybrid model by linking the X-ray structure of crTP_part2_RnaH_ to the AlphaFold-predicted structure of RnaH and fitted against the experimental data. The *p(r)* function and the dimensionless Kratky plot showed an elongated particle (**Fig. S7**). The EOM analysis revealed that there are three distinct conformations present in solution for crTP_part2_RnaH_-RnaH (**Fig. 2C**) indicating flexibility between the crTP_part2_RnaH_ domain and the attached cargo protein.

In contrast, crTP_part2_ArgC_-ArgC appears to be dimeric over the whole concentration range (max. 10 mg/ml). An AlphaFold2 model prediction of the dimer showed the dimer interface within the cargo protein ArgC. The *p(r)* function and the dimensionless Kratky plot showed elongated multidomain particle (**Fig. S8**). Similar to crTP_part2_RnaH_-RnaH, we created a hybrid model using the ArgC dimer interface conformation from AlphaFold as rigid body and allowed the crTP_part2_ArgC_ part from the solved crystal structure to move in the EOM analysis ((χ^2^ of 1.179) (**Fig. 2D**, **Table S2**). The models obtained showed that crTP_part2_ArgC_-ArgC forms an overall compact molecule but, reveals flexibility between the crTP_part2_ domain and the attached cargo protein (**Fig. 2D** and **Fig. S8**). Furthermore, the N-terminal tail of crTP_part2_ArgC_ remained flexible.

### **γ**-Glutamyl cyclotransferase fold proteins

For several proteins in the GGCT-like superfamily, enzymatic functions have been characterized. GGCTs degrade diverse γ-glutamyl-dipeptides to 5-oxo-proline and a free amino acid. The γ-glutamylamine-cyclotransferase (GGACT) family is specialized for the breakdown of γ-glutamyl-ε-lysine into 5-oxo-proline and lysine. Whereas ChaC proteins preferentially catalyze the breakdown of γ-glutamyl-cysteinyl-glycine (i.e., glutathione) to 5- oxo-proline and Cys-Gly. The BtrG protein from *Bacillus circulans* (= *Niallia circulans*) that is the name-giving protein of the prokaryotic BtrG-like family removes by cyclotransferase activity a protective γ-glutamyl group that is required during the biosynthesis of the antibiotic butirosin (Llewellyn et al., 2007). Thus, a common theme for all these enzymes is the recognition and cleavage of diverse γ-glutamyl derivatives. Finally, AIG2A and AIG2B of the plant-specific AIG2/AIG2-like family, recently have been characterized as immune regulators in *Arabidopsis thaliana* that balance different chemical defense systems but their exact molecular mode of action is unclear (Wang et al., 2022).

## Supplemental Methods

### Construction of expression plasmids

For the construction of expression vectors, initially, the sequence coding for crTP_part2_RnaH_ was amplified from cDNA using primer pair 1515+1514 and cloned into pTEV-16b, a vector based on pET-16b (Novagene, 69662) in which the sequence for a TEV cleavage site was introduced after the sequence for the N-terminal His_10_- tag by Gibson cloning (Gibson et al., 2009) resulting in vector GPN125. As a backbone, for the expression vectors used in this study, vector GPN131 was used. This vector is a derivative of the plasmid pET-22b(+) (Novagene, 69744), in which the sequence for the pelB leader and C-terminal His_6_-tag were replaced by the sequence for an N-terminal His_6_-tag, thrombin cleavage site, and SUMO-tag from the plasmid pET-6His-SUMO-MarathonRT (NovoPro, V014422) by Golden Gate cloning (Engler et al., 2008). The His_6_-SUMO-TEV-crTP_part2_RnaH_ expression vector (GPN142) was generated by amplifying the sequence coding for the TEV cleavage site linked to the N-terminus of crTP_part2_RnaH_ using primer pairs 1782+1783 (binding to the insert on template vector GPN125) and 1785+1784 (binding to the backbone vector GPN131), resulting ultimately in a pET22b(+) derivative carrying the insert shown in **Fig. S10**. All other expression vectors were generated by replacing or inserting sequences amplified from *P. chromatophora* cDNA into GPN142 or derivatives thereof by Gibson cloning. To generate the His_6_-SUMO-TEV-crTP_part2_RnaH_-RnaH expression vector (GPN194), the RnaH-coding sequence was inserted into GPN142 downstream of the crTP_part2_RnaH_-coding sequence using fragments amplified with the primer pairs 2640+2641 (binding to cDNA derived from scaffold2581) and 2639+2638 (binding to the backbone vector GPN142). To generate the His_6_- SUMO-TEV-crTP_part2_ArgC_ expression vector (GPN167), the crTP_part2_RnaH_-coding sequence in GPN142 was replaced by the crTP_part2_ArgC_-coding sequence using fragments amplified with the primer pairs 2297+2298 (binding to cDNA derived from scaffold7023) and 2299+2300 (binding to the backbone vector GPN142). To generate the His_6_-SUMO-TEV-crTP_part2_ArgC_- ArgC expression vector (GPN195), the ArgC-coding sequence was inserted into GPN167 using fragments amplified with the primer pairs 2669+2645 (binding to from cDNA derived from scaffold7023) and 2643+2670 (binding to the backbone vector GPN167). To generate the His_6_- SUMO-TEV-crTP_part2_CysK_ expression vector (GPN168), the crTP_part2_RNAH_-coding sequence in GPN142 was replaced by the crTP_part2_CysK_-coding sequence using fragments amplified with the primer pairs 2293+2294 (binding to cDNA derived from scaffold4337) and 2295+2296 (binding to the backbone vector GPN142). All primer sequences are provided in **Table S5**. For cloning procedures and plasmid propagation, *E. coli* Top10 (Invitrogen, Carlsbad, CA, USA) was used. The correct nucleotide sequence of all constructs was confirmed by sequencing.

### Conditions for heterologous expression of recombinant proteins

Expression cultures in Luria Bertani (LB) (Sambrook et al., 1998) or 2YT medium containing 100 µg/ml ampicillin and 34 µg/ml chloramphenicol were inoculated to an OD_600_ of 0.1 and grown at 37°C and 180 rpm to an OD_600_ of 0.6. Then, protein expression was induced by 0.5 mM isopropyl-β-D- thiogalactopyranoside (IPTG) under the following conditions optimized for maximal yield. Cells containing GPN125, GPN167, GPN194, and GPN195 were grown in 2 L 2YT medium, and shifted to 16°C before induction for 24 h. Cells containing GPN168 were grown in 2 L LB medium and maintained at 37°C for 3 h after induction. Finally, cells were harvested by centrifugation (5,000 × g for 15 min).

### Protein purification

Frozen cells from expression cultures were resuspended in 20 mM HEPES (pH 8.0) and 300 mM NaCl (buffer A) supplemented with DNaseI (Roche, product no. 1010415900) and 1-fold concentrated protease inhibitor (cOmplete, EDTA-free, Roche). Cells were disrupted by sonication (Hielscher, UP50H; equipped with the MS3 probe) using pulse-pause cycles of 30 s at an amplitude of 60% for 16 minutes. Throughout the process, samples were cooled to 4°C. Cell debris and insoluble material were removed in two steps by centrifugation at 12,000 × g for 15 min and then at 120,000 × g for 1 h (45Ti fixed-angle rotor, Beckman Coulter). The supernatant was applied to 5 ml Ni-NTA columns (HisTrap HP 5 ml, Cytiva) equilibrated with buffer A in an ÄKTA Pure 25 system (Cytiva, formerly GE Healthcare) following the manufacturer’s recommendations. The column was washed with 10 column volumes (CVs) of buffer A, followed by a linear gradient of 20 CVs targeting 100 % buffer A + 500 mM imidazole. All five recombinant proteins showed a similar elution pattern with two peaks in λ = 280 nm absorption measurements. To avoid contamination from the first peak, fractions resulting from the second half of the second peak were collected and pooled. This corresponded to 120-210 mM imidazole for His_6_-SUMO-TEV-crTP_part2_RnaH,_ 135-300 mM imidazole for His_6_-SUMO-TEV-crTP_part2_ArgC,_ 140-190 mM imidazole for His_6_-SUMO-TEV- crTP_part2_CysK_, and 125-250 mM imidazole for His_6_-SUMO-TEV-crTP_part2ArgC_-ArgC, and 90-200 mM imidazole for His6-SUMO-TEV-crTPpart2_RnaH-RnaH. Elution fractions were diluted 1:2 with buffer A to decrease imidazole concentration and the His_6_-SUMO tag was cleaved off with TEV-protease in a molar ratio of 1:100 (rotating at 4°C overnight). The cleaved tags were removed by reverse IMAC on 5 ml Ni-NTA columns (HisTrap HP 5 ml, Cytiva). The flow-through was collected and concentrated to 5 ml (Amicon® Ultra Centrifugal Filters, 10 kDa MWCO). Residual contaminants in this concentrated flow-through were removed by size exclusion chromatography (SEC; Superdex® 200 Increase 10/300 GL, Cytiva). The entire process was carried out at 4°C, at a set maximal flow rate of 0.8 ml/min.

### Protein crystallization and 3D structure determination by X-ray crystallography

crTP_part2_RnaH_ crystals formed after 5 d. For experimental phasing, the crystals were soaked with 0.3 µl 10 mM potassium hexachloroiridate (IV). The crystals were cryo-protected by adding 1 µl ethylene glycol to the drop and flash frozen. crTP_part2_ArgC_ crystals formed after 41 d. The crystals were cryo-protected by adding mineral oil over the drop, dragged through it and flash frozen.

For all collected datasets, data reduction was performed using XDS (Kabsch, 2010) and aimless (Evans and Murshudov, 2013) from the CCP4 suite (Winn et al., 2011). The structure of crTP_part2_RnaH_ was initially phased using ShelX (Usón and Sheldrick, 2018) which is part of the CCP4 v8.0 suit and after manually building the backbone structure, this model was used to solve the high resolution data set via molecular replacement with Phaser (McCoy et al., 2007). The initial model was refined alternating cycles of manual model building in COOT (Emsley and Cowtan, 2004; Emsley et al., 2010) and automatic refinement using Phenix (Liebschner et al., 2019) version 1.19.2_4158. Data collection and refinement statistics are reported in Table S1. The final model of crTP_part2_RnaH_ was then used as template for molecular replacement and phasing of the crTP_part2_ArgC_ dataset.

### SAXS

The sample-to-detector distance of the P12 beamline was 3.00 m, resulting in an achievable q-range of 0.03-7.0 nm^-1^. Measurements were performed in batch mode at 10°C with a protein concentration range of 9.30-1.16 mg/ml for crTP_part2_RnaH_, 8.6-0.6 mg/ml for crTP_part2_ArgC_, and 10.6-0.7 mg/ml for crTP_part2_CysK_. We collected 40 frames per protein sample. Exposure time was 0.095 s/frame. Data were scaled to absolute intensity against water. We extrapolated the datasets of crTP_part2_ArgC_ and crTP_part2_CysK_ to zero concentration and merged crTP_part2_RnaH_ data from high and low concentrations.

On our Xeuss 2.0 Q-Xoom system (equipped with a PILATUS 3 R 300K detector (Dectris) and a GENIX 3D CU Ultra Low Divergence x-ray beam delivery system), the chosen sample-to-detector distance was 0.55 m, resulting in an achievable q-range of 0.05-6 nm^-1^. The measurement was performed at 10°C with a protein concentration range of 0.62-10.00 mg/ml for crTP_part2_ArgC_-ArgC and 1.00-9.00 mg/ml for, crTP_part2_RnaH_-RnaH respectively. The samples were injected in the Low Noise Flow Cell (Xenocs) via autosampler. We collect 18 frames (exposure time: 10 min/frame) and scaled the data to absolute intensity against water. We checked the data for concentration effect and used the 10.00 mg/ml concentration data for crTP_part2_ArgC_-ArgC and 9.00 mg/ml data of crTP_part2_RnaH_-RnaH for further analysis. All used programs for data processing were part of the ATSAS Software package (Version 3.0.5) (Manalastas-Cantos et al., 2021).

For the remodeling of the flexible N and C terminal parts of solved crTP_part2_RnaH_ we used EOM (Bernadó et al., 2007; Tria et al., 2015). Some terminal residues were missing in the crystal structure of crTP_part2_ArgC_ so we added these residues with CORAL (Petoukhov et al., 2012) to complete the solved crystal structure. We used the solved part of the crystal structure from crTP_part2_ArgC_ and created a homology model of crTP_part2_CysK_ with SWISS-MODEL (Waterhouse et al., 2018) and completed and remodel the tails with CORAL (Petoukhov et al., 2012). We created hybrid models of crTP_part2_ArgC_-ArgC and crTP_part2_RnaH_-RnaH, based on the corresponding solved crTP_part2_ parts and the cargo proteins predictions from AlphaFold2 (Jumper et al., 2021; Mirdita et al., 2022) and remodeled the flexible parts with EOM.

---

**Figure S1:**
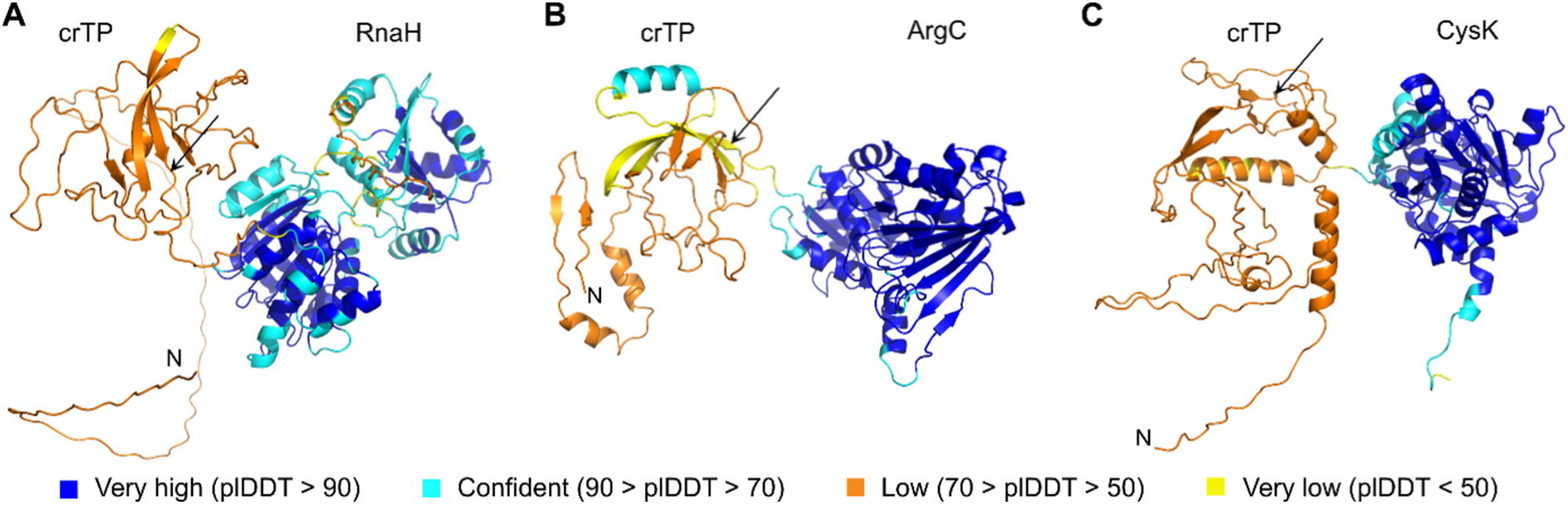
AlphaFold3-perdiceted protein models. Proteins encoded by transcripts scaffold2581 **(A),** scaffold7023 **(B)**, and scaffold7023 **(C)** were analyzed with AlphaFold3. N-termini (N) are indicated and the start of the linker between the crTP and the cargo protein is highlighted by a black arrow. The confidence estimate (pLDDT) is shown as color code in the images of the structures.

**Figure S2:**
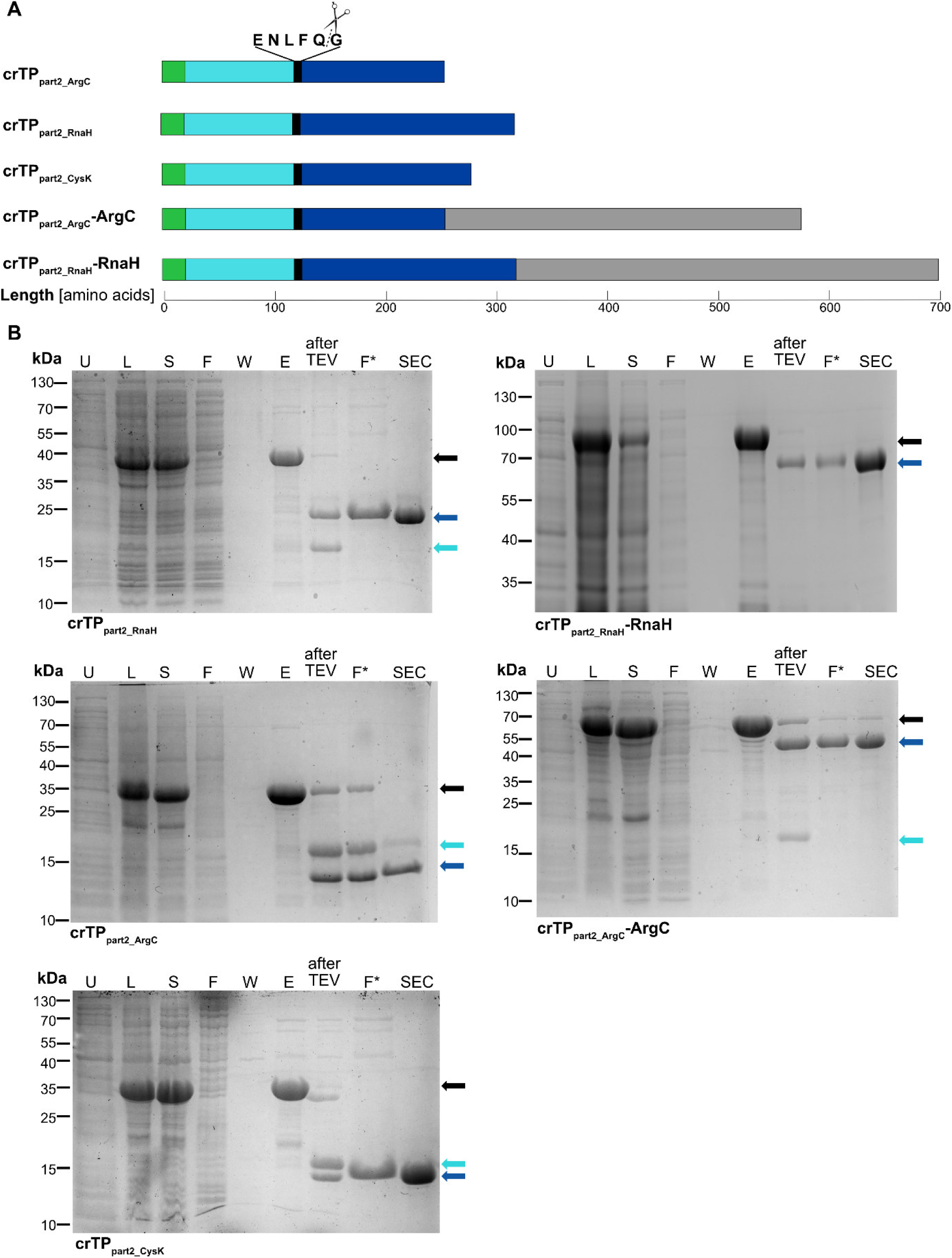
**Purification of crTPpart2-containing constructs. (A)** Schematic representation of crTPpart2 constructs. The His6-tag (green) and SUMO solubility-tag (cyan) are cleaved off at the TEV protease recognition site (black) to obtain crTPpart2 (blue) alone or attached to its corresponding cargo protein (grey). **(B)** All crTPpart2 constructs were expressed in *E. coli* following induction with IPTG. For un- induced samples (U), 600 µl of expression culture was withdrawn before induction, spun down, and the pellet resuspended in 60 µl PBS. Following expression, lysate (L) was generated spun at 120,000 x g, for 1 h. The supernatant (S) was loaded onto a Ni-NTA column. Column was washed with buffer A. Samples from the flow-through (F) and mid-wash (W) were collected. Proteins of interest were eluted (E). Eluate was diluted 1:2 with buffer A and digested with TEV-protease (‘after TEV’). Proteins of interest were isolated by reverse IMAC by collecting the flow-through (F*). Then, proteins of interest were further purified by SEC. Proteins in all samples were solubilized in Laemmli buffer and 5 µl loaded onto a 12.5 % polyacrylamid gel or 10 % for crTPpart2_RnaH-RnaH. For F* and after SEC, ∼5 µg total protein was loaded on the gel. The gel was stained with Coomassie brilliant blue. Full-length constructs are indicated by black arrows, proteins of interest by blue arrows, and His-SUMO (14.2 kDa) by cyan arrows.

**Figure S3:**
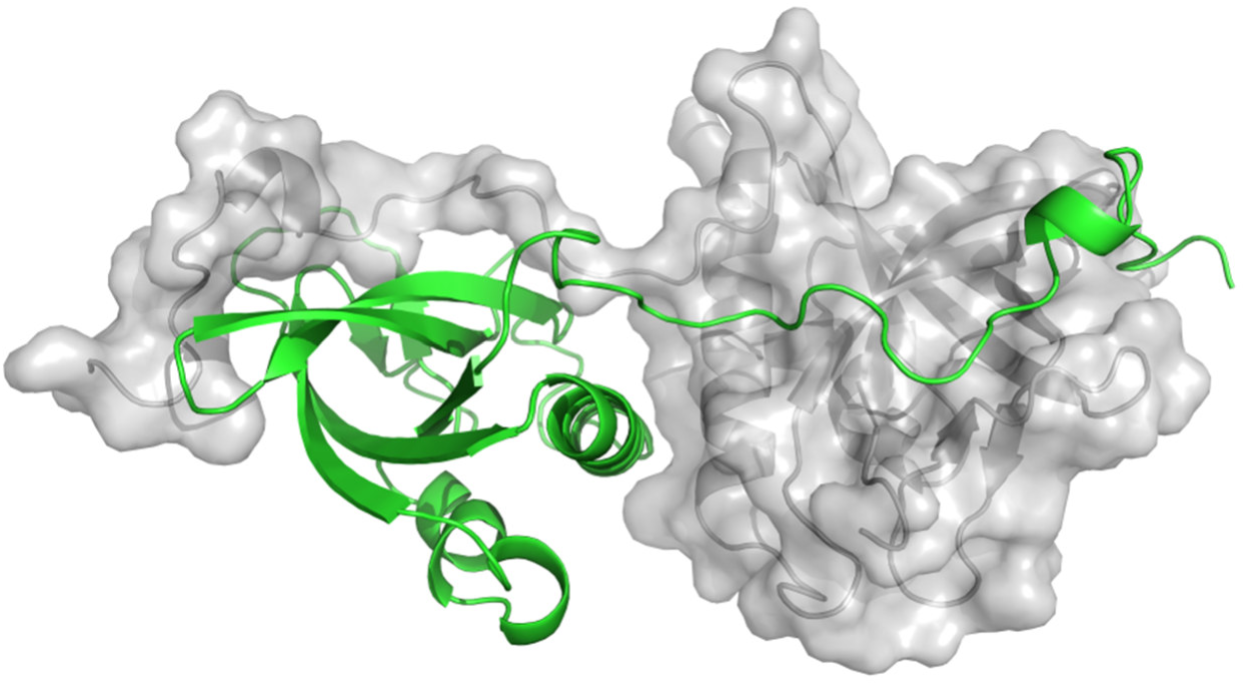
Crystallographic dimer of crTPpart2_RnaH. One monomer is shown as green cartoon model, the other as grey surface structure.

**Figure S4:**
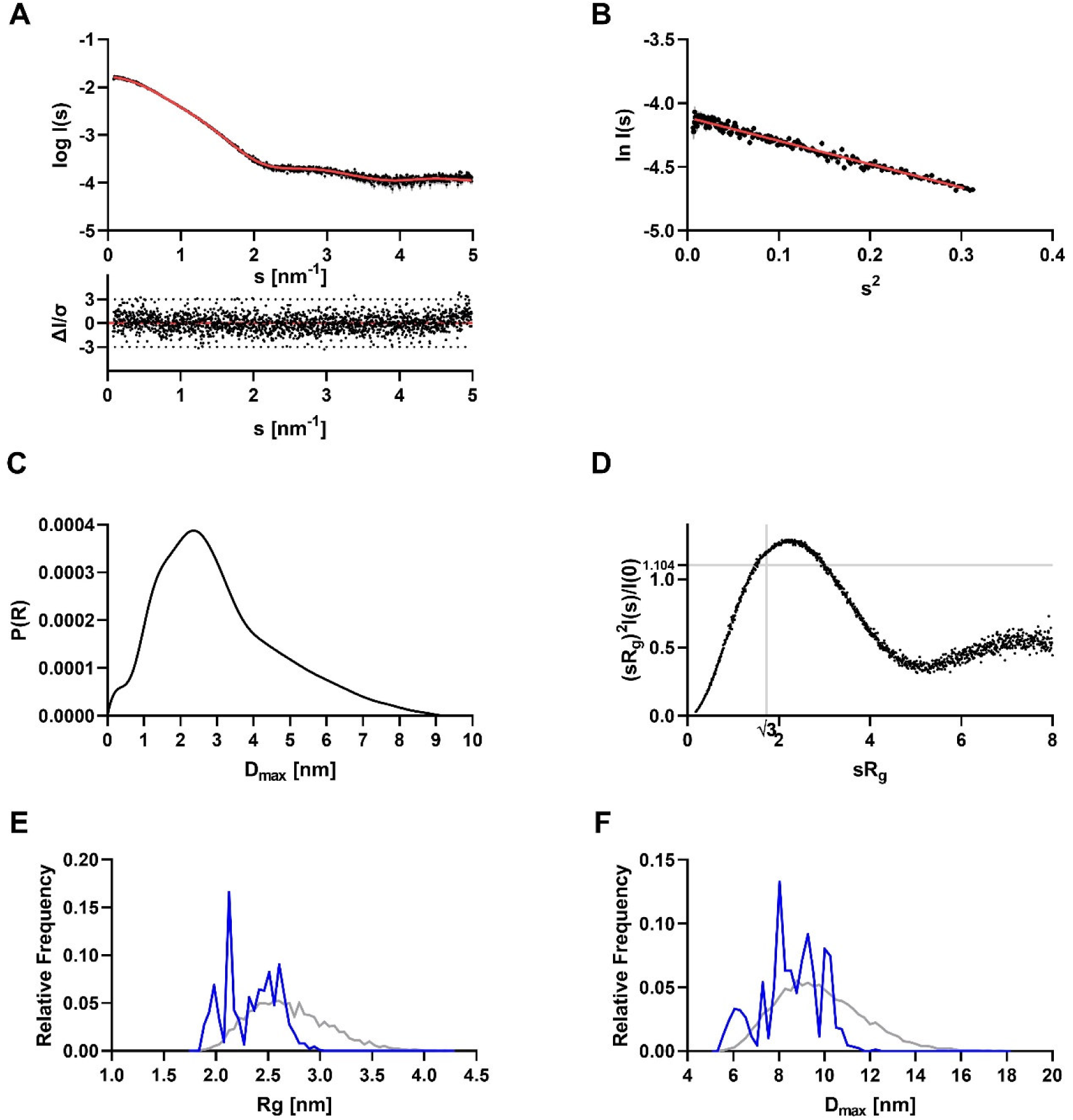
**Small-angle X-ray scattering data from crTPpart2_RnaH. A:** Scattering data of crTPpart2_RnaH. Experimental data are shown in black dots, with grey error bars. The EOM ensemble model fit is shown as red line and below is the residual plot of the data. **B:** The Guinier plot of crTPpart2_RnaH showed a stable Guinier region with a *Rg* of 2.34 nm. **C:** The *p(r)* function of crTPpart2_RnaH showed an elongated particle with a *Dmax* value of 9.17 nm. **D:** The dimensionless Kratky plot of crTPpart2_RnaH showed an elongated particle with a degree of flexibility of the termini. **E, F:** *Rg* and *Dmax* distribution of crTPpart2_RnaH. Ensemble pool is shown in grey, selected EOM models are shown in blue.

**Figure S5:**
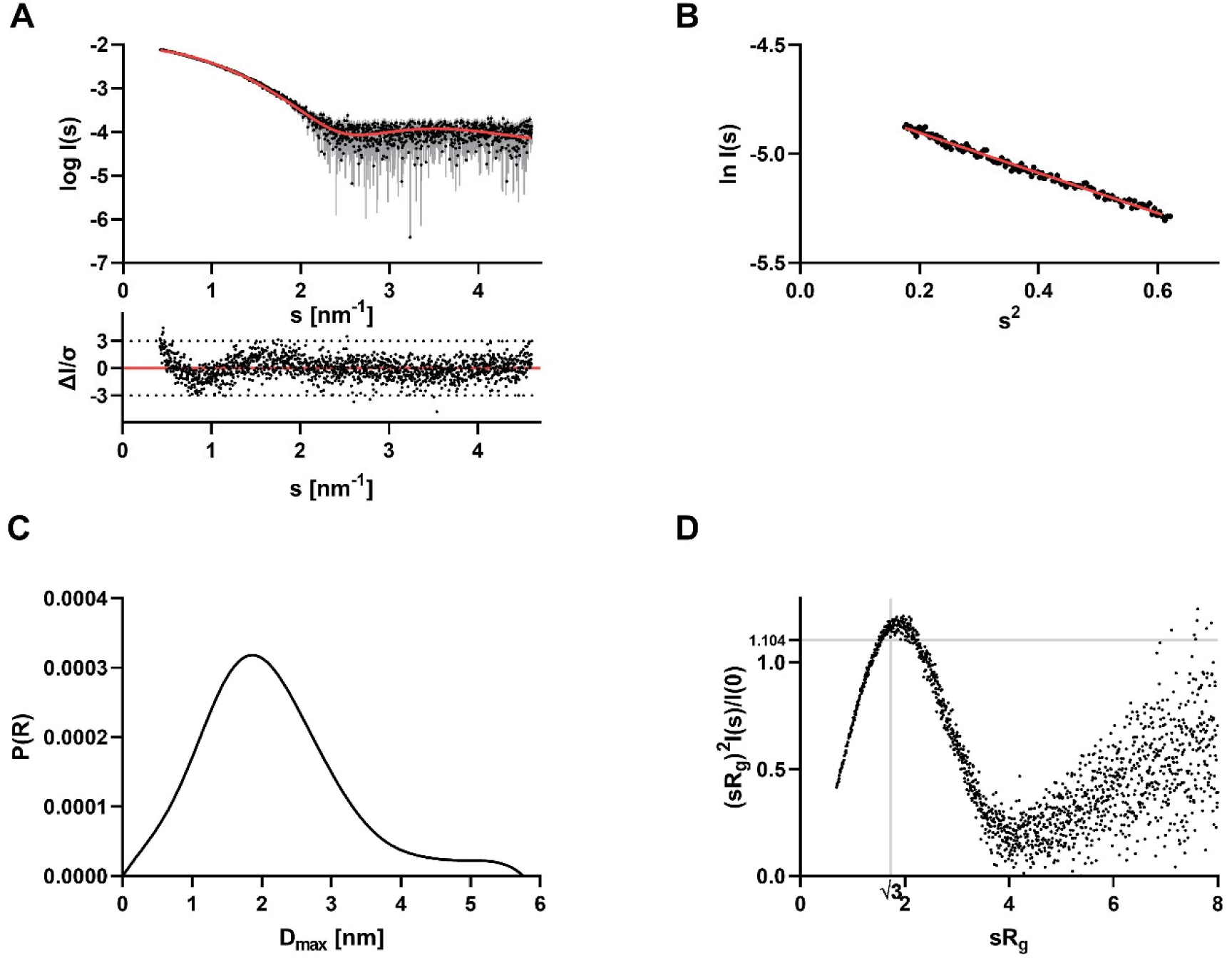
**Small-angle X-ray scattering data from crTPpart2_ArgC. A:** Scattering data of crTPpart2_ArgC. Experimental data are shown in black dots, with grey error bars. The CORAL model fit is shown as red line and below is the residual plot of the data. **B:** The Guinier plot of crTPpart2_ArgC showed a stable Guinier region with a *Rg* of 1.66 nm. **C:** The *p(r)* function of crTPpart2_ArgC showed a globular molecule with an elongated part and a *Dmax* value of 5.76 nm. **D:** The dimensionless Kratky plot of crTPpart2_ArgC showed a little elongated, but compact molecule.

**Figure S6:**
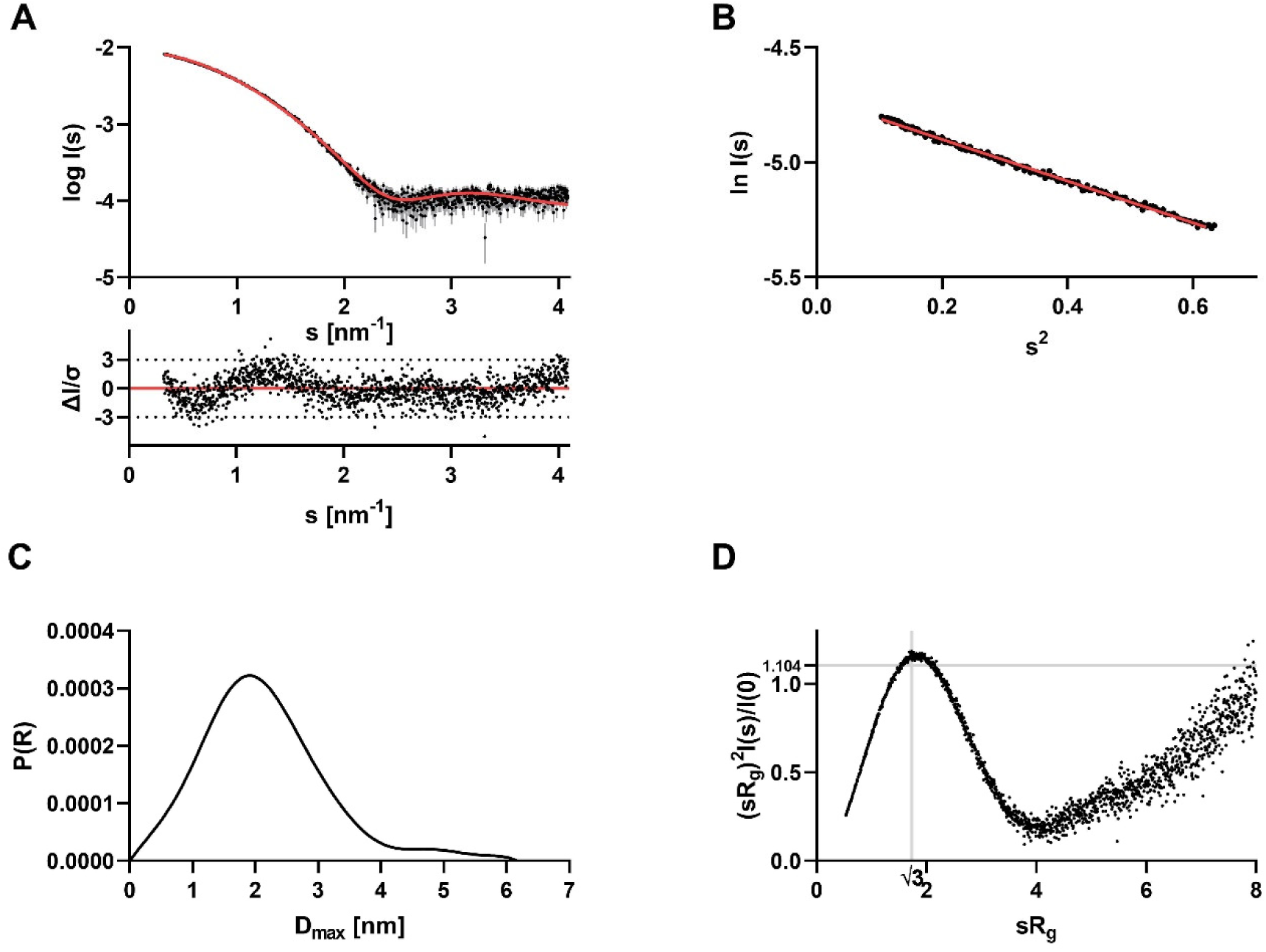
**Small-angle X-ray scattering data from crTPpart2_CysK. A:** Scattering data of crTPpart2_CysK. Experimental data are shown in black dots, with grey error bars. The CORAL model fit is shown as red line and below is the residual plot of the data. **B:** The Guinier plot of crTPpart2_CysK showed a stable Guinier region with a *Rg* of 1.65 nm. **C:** The *p(r)* function of crTPpart2_CysK showed a globular molecule with an elongated part and a *Dmax* value of 6.17 nm. **D:** The dimensionless Kratky plot of crTPpart2_CysK showed a little elongated, but compact molecule.

**Figure S7:**
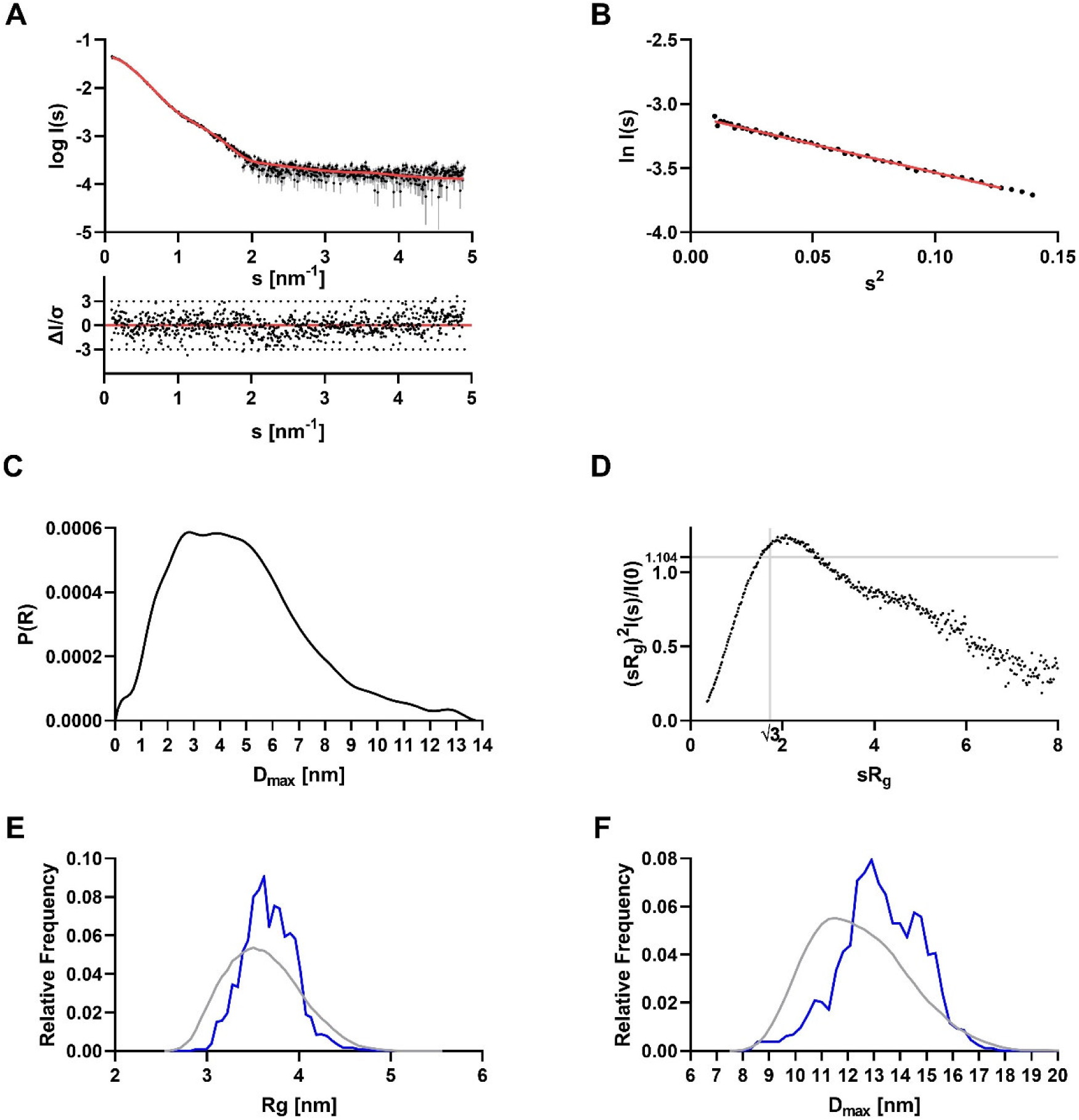
**Small-angle X-ray scattering data from crTPpart2_RnaH-RnaH. A:** Scattering data of crTPpart2_RnaH-RnaH. Experimental data are shown in black dots, with grey error bars. The EOM ensemble model fit is shown as red line and below is the residual plot of the data. **B:** The Guinier plot of crTPpart2_RnaH-RnaH showed a stable Guinier region with a *Rg* of 3.65 nm. **C:** The *p(r)* function of crTPpart2_RnaH-RnaH showed an elongated multidomain particle with a *Dmax* value of 13.73 nm. **D:** The dimensionless Kratky plot of crTPpart2_RnaH-RnaH showed an elongated multidomain particle. **E and F:** *Rg* and *Dmax* distribution of crTPpart2_RnaH-RnaH. Ensemble pool is shown in grey, selected EOM models are shown in blue.

**Figure S8:**
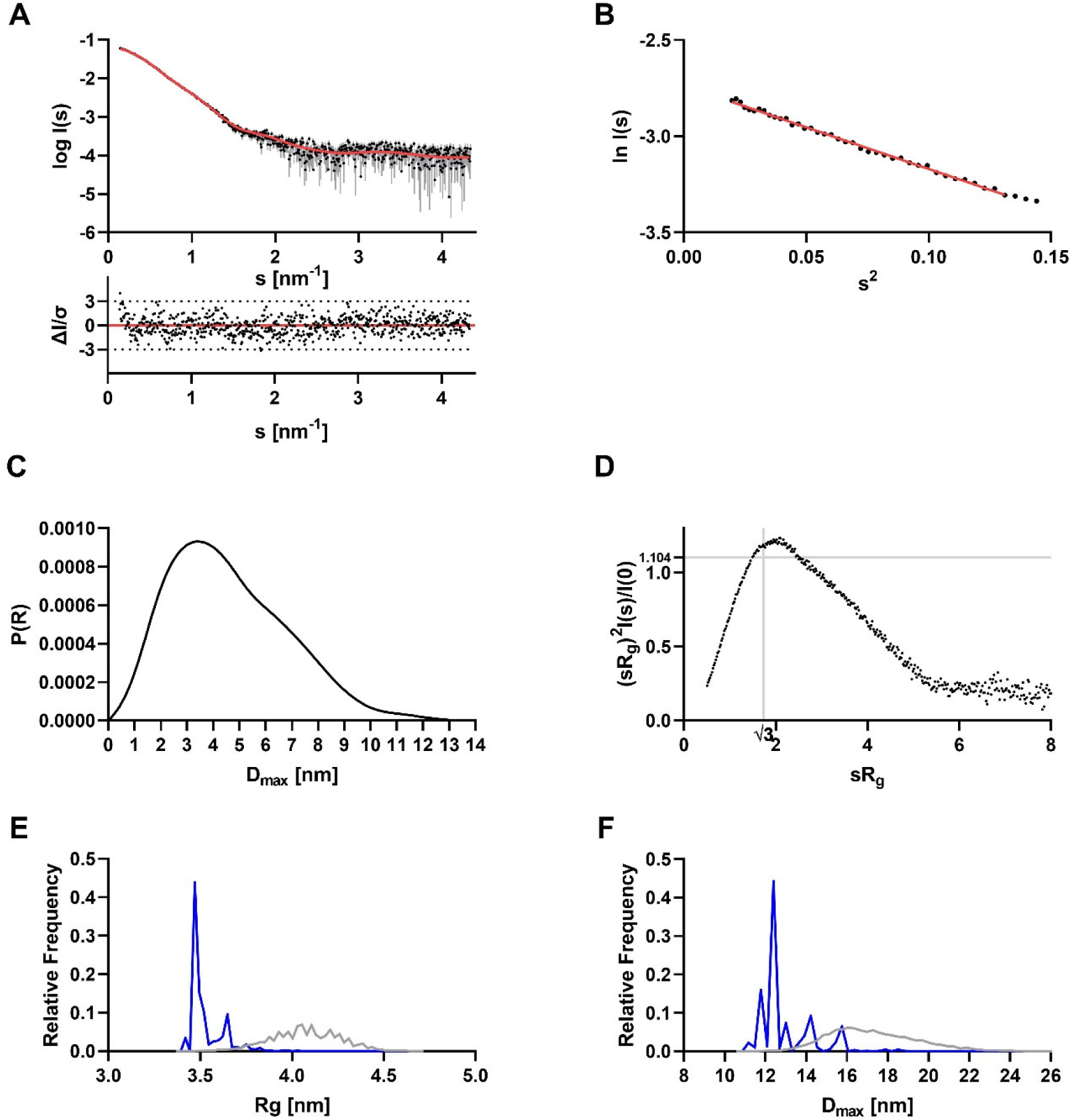
**Small-angle X-ray scattering data from crTPpart2_ArgC-ArgC. A:** Scattering data of crTPpart2_ArgC-ArgC. Experimental data are shown in black dots, with grey error bars. The EOM ensemble model fit is shown as red line and below is the residual plot of the data. **B:** The Guinier plot of crTPpart2_ArgC-ArgC showed a stable Guinier region with a *Rg* of 3.60 nm. **C:** The *p(r)* function of crTPpart2_ArgC-ArgC showed an elongated multidomain particle with a *Dmax* value of 13.11 nm. **D:** The dimensionless Kratky plot of crTPpart2_ArgC-ArgC showed an elongated multidomain particle. **E &F:** *Rg* and *Dmax* distribution of crTPpart2_ArgC-ArgC. Ensemble pool is shown in grey, selected EOM models are shown in blue.

**Figure S9:**
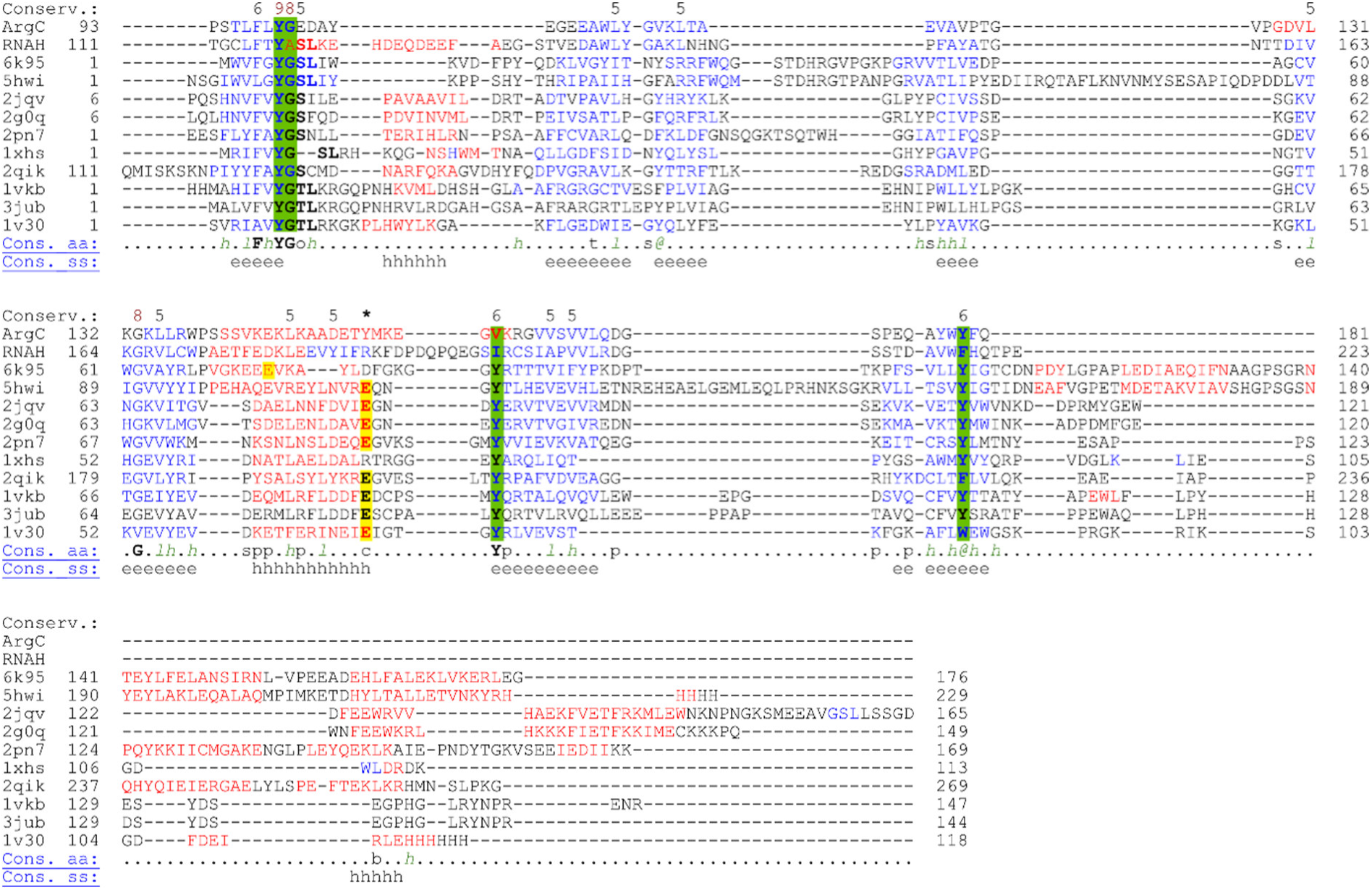
**The typical GGCT binding pocket is conserved but the active site is not in crTPpart2 structures.** Structure-based alignment of GGCT-like proteins generated with Promals3D (Pei et al., 2008) using default parameters. Secondary structure predictions according to PSIPRED (Jones, 1999): red, α-helix; blue, β-strand. Consensus sequence is provided if the weighted frequency of a certain class of residues in a position is above 0.8. Here, conserved aas are in bold uppercase letters; aliphatic (I, V, L): l; aromatic (Y, H, W, F): @; hydrophobic (W, F, Y, M, L, I, V, A, C, T, H): h; alcohol (S, T): o; polar residues (D, E, H, K, N, Q, R, S, T): p; tiny (A, G, C, S): t; small (A, G, C, S, V, N, D, T, P): s; bulky residues (E, F, I, K, L, M, Q, R, W, Y): b; positively charged (K, R, H): +; negatively charged (D, E): -; charged (D, E, K, R, H): c. Note that in the human ChaC2 (pdb id: 6k95) the catalytic site (Glu74) moved to a long flexible loop. By dimerization of two ChaC2 monomers, the flexible loop of one monomer moves into the catalytic cavity of the other monomer resulting in Glu74 to come close in position to the catalytic centers of other GGCT-like proteins (Nguyen et al., 2020). In the *E. coli* homolog of BtrG (named YftP, of unknown function; pdb id 1xhs) Glu82 of 3jub is replaced by an Arg which suggests that the protein does not have cyclotransferase activity.

**Figure S10:**
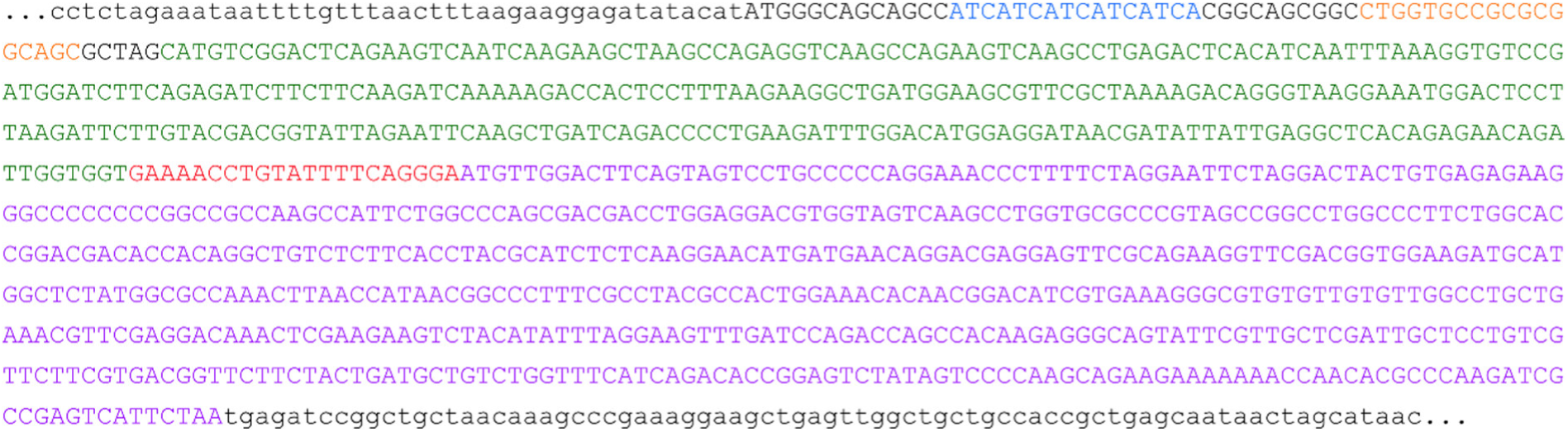
**The GPN142 expression vector is a pET22b(+) derivative, carrying the depicted insert**. Into the backbone pET22b(+) backbone (lowercase letters) an insert containing the coding sequences for His6 (blue), a thrombin cleavage site (orange), a SUMO-tag (green), a TEV cleavage site (red), and the crTPpart2_RnaH domain (violet) was inserted at the indicated position.

**Table S1:** X-ray crystallography data collection and refinement statistics for crTP_part2_RnaH_ and crTP_part2_ArgC_. Statistics for the highest resolution shell are shown in parentheses.

|  | crTP<sub>part2_RnaH</sub> (PDB ID: 9I09) | crTP<sub>part2_ArgC</sub> (PDB ID: 9I08) |
| --- | --- | --- |
| **Wavelength** | 1.000 | 0.9763 |
| **Resolution range** | 35.15 - 2.4 (2.49 - 2.4) | 43.75 - 2.2 (2.28-2.2) |
| **Space group** | P 31 2 1 | P 1 21 1 |
| **Unit cell** | 81.164 81.164 63.678 90 90 120 | 36.069 67.227 43.774 90 92.03 90 |
| **Total reflections** | 19658 (1874) | 20934 (1855) |
| **Unique reflections** | 9829 (937) | 10544 (944) |
| **Multiplicity** | 2.0 (2.0) | 2.0 (2.0) |
| **Completeness (%)** | 100 (99.9) | 98.5 (87.6) |
| **Mean I/sigma(I)** | 34.0 (11.4) | 15.6 (6.9) |
| **Wilson B-factor** | 46.61 | 22.48 |
| **R-merge** | 0.009 (0.043) | 0.03 (0.162) |
| **R-meas** | 0.013 (0.06) | 0.042 (0.162) |
| **R-pim** | 0.009 (0.043) | 0.03 (0.115) |
| **CC1/2** | 1.00 (0.997) | 0.998 (0.97) |
| **Reflections used in refinement** | 18682 (820) | 19417 (1710) |
| **Reflections used for R-free** | 976 (117) | 977 (145) |
| **R-work** | 0.1860 (0.2017) | 0.2034 (0.2685) |
| **R-free** | 0.2203 (0.2676) | 0.2260 (0.3033) |
| **Number of non-hydrogen atoms** | 1223 | 2235 |
| **macromolecules** | 1157 | 1910 |
| **ligands** | 4 | 16 |
| **solvent** | 62 | 309 |
| **Protein residues** | 146 | 247 |
| **RMS(bonds)** | 0.007 | 0.009 |
| **RMS(angles)** | 0.95 | 0.81 |
| **Ramachandran favored (%)** | 96.53 | 96.71 |
| **Ramachandran allowed (%)** | 3.47 | 2.47 |
| **Ramachandran outliers (%)** | 0.00 | 0.82 |
| **Rotamer outliers (%)** | 0.81 | 0.98 |
| **Clashscore** | 4.44 | 7.25 |
| **Average B-factor** | 52.96 | 26.28 |
| **macromolecules** | 53.00 | 25.44 |
| **ligands** | 41.42 | 38.48 |
| **solvent** | 52.80 | 30.85 |

**Table S2:**
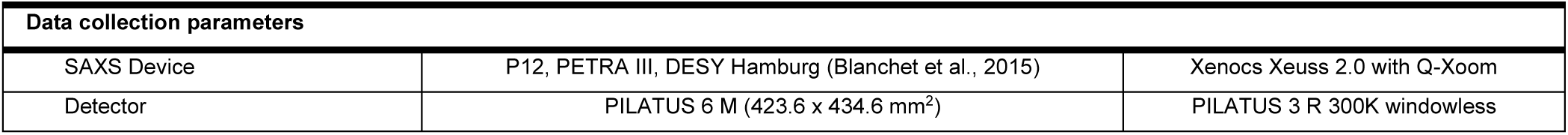
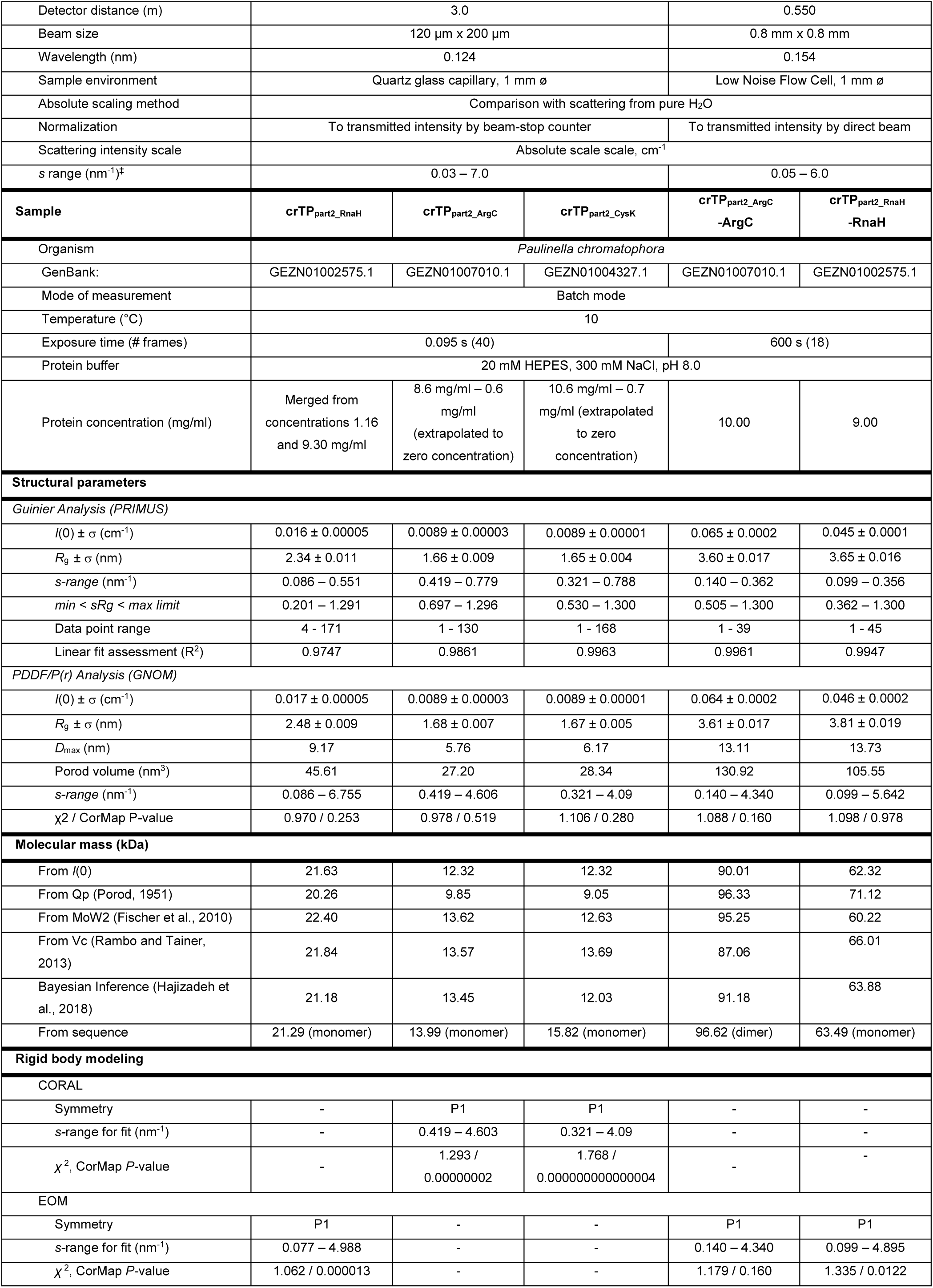
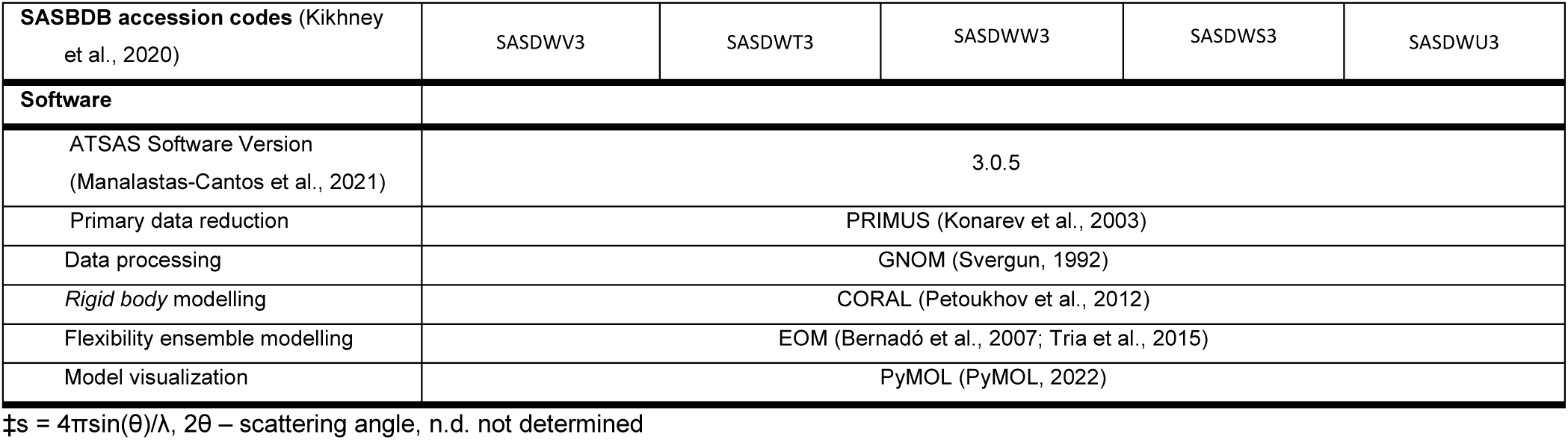
Overall SAXS Data.

| Data collection parameters |  |  |
| --- | --- | --- |
| SAXS Device | P12, PETRA III, DESY Hamburg (Blanchet et al., 2015) | Xenocs Xeuss 2.0 with Q-Xoom |
| Detector | PILATUS 6 M (423.6 x 434.6 mm<sup>2</sup>) | PILATUS 3 R 300K windowless |

|  |  |  |  |  |  |
| --- | --- | --- | --- | --- | --- |
| Detector distance (m) | 3.0 |  |  | 0.550 |  |
| Beam size | 120 μm x 200 μm |  |  | 0.8 mm x 0.8 mm |  |
| Wavelength (nm) | 0.124 |  |  | 0.154 |  |
| Sample environment | Quartz glass capillary, 1 mm ø |  |  | Low Noise Flow Cell, 1 mm ø |  |
| Absolute scaling method | Comparison with scattering from pure H2O |  |  |  |  |
| Normalization | To transmitted intensity by beam-stop counter |  |  | To transmitted intensity by direct beam |  |
| Scattering intensity scale | Absolute scale scale, cm<sup>-1</sup> |  |  |  |  |
| s range (nm<sup>-1</sup>)<sup>‡</sup> | 0.03 – 7.0 |  |  | 0.05 – 6.0 |  |
| Sample | crTP<sub>part2_RnaH</sub> | crTP<sub>part2_ArgC</sub> | crTP<sub>part2_CysK</sub> | crTP<sub>part2_ArgC</sub><br>-ArgC | crTP<sub>part2_RnaH</sub><br>-RnaH |
| Organism | Paulinella chromatophora |  |  |  |  |
| GenBank: | GEZN01002575.1 | GEZN01007010.1 | GEZN01004327.1 | GEZN01007010.1 | GEZN01002575.1 |
| Mode of measurement | Batch mode |  |  |  |  |
| Temperature (°C) | 10 |  |  |  |  |
| Exposure time (# frames) | 0.095 s (40) |  |  | 600 s (18) |  |
| Protein buffer | 20 mM HEPES, 300 mM NaCl, pH 8.0 |  |  |  |  |
| Protein concentration (mg/ml) | Merged from concentrations 1.16 and 9.30 mg/ml | 8.6 mg/ml – 0.6 mg/ml (extrapolated to zero concentration) | 10.6 mg/ml – 0.7 mg/ml (extrapolated to zero concentration) | 10.00 | 9.00 |
| Structural parameters |  |  |  |  |  |
| Guinier Analysis (PRIMUS) |  |  |  |  |  |
| I(0) ± σ (cm<sup>-1</sup>) | 0.016 ± 0.00005 | 0.0089 ± 0.00003 | 0.0089 ± 0.00001 | 0.065 ± 0.0002 | 0.045 ± 0.0001 |
| R<sub>g</sub> ± σ (nm) | 2.34 ± 0.011 | 1.66 ± 0.009 | 1.65 ± 0.004 | 3.60 ± 0.017 | 3.65 ± 0.016 |
| s-range (nm<sup>-1</sup>) | 0.086 – 0.551 | 0.419 – 0.779 | 0.321 – 0.788 | 0.140 – 0.362 | 0.099 – 0.356 |
| min < sR<sub>g</sub> < max limit | 0.201 – 1.291 | 0.697 – 1.296 | 0.530 – 1.300 | 0.505 – 1.300 | 0.362 – 1.300 |
| Data point range | 4 - 171 | 1 - 130 | 1 - 168 | 1 - 39 | 1 - 45 |
| Linear fit assessment (R<sup>2</sup>) | 0.9747 | 0.9861 | 0.9963 | 0.9961 | 0.9947 |
| PDDF/P(r) Analysis (GNOM) |  |  |  |  |  |
| I(0) ± σ (cm<sup>-1</sup>) | 0.017 ± 0.00005 | 0.0089 ± 0.00003 | 0.0089 ± 0.00001 | 0.064 ± 0.0002 | 0.046 ± 0.0002 |
| R<sub>g</sub> ± σ (nm) | 2.48 ± 0.009 | 1.68 ± 0.007 | 1.67 ± 0.005 | 3.61 ± 0.017 | 3.81 ± 0.019 |
| D<sub>max</sub> (nm) | 9.17 | 5.76 | 6.17 | 13.11 | 13.73 |
| Porod volume (nm<sup>3</sup>) | 45.61 | 27.20 | 28.34 | 130.92 | 105.55 |
| s-range (nm<sup>-1</sup>) | 0.086 – 6.755 | 0.419 – 4.606 | 0.321 – 4.09 | 0.140 – 4.340 | 0.099 – 5.642 |
| χ<sup>2</sup> / CorMap P-value | 0.970 / 0.253 | 0.978 / 0.519 | 1.106 / 0.280 | 1.088 / 0.160 | 1.098 / 0.978 |
| Molecular mass (kDa) |  |  |  |  |  |
| From I(0) | 21.63 | 12.32 | 12.32 | 90.01 | 62.32 |
| From Qp (Porod, 1951) | 20.26 | 9.85 | 9.05 | 96.33 | 71.12 |
| From MoW2 (Fischer et al., 2010) | 22.40 | 13.62 | 12.63 | 95.25 | 60.22 |
| From Vc (Rambo and Tainer, 2013) | 21.84 | 13.57 | 13.69 | 87.06 | 66.01 |
| Bayesian Inference (Hajizadeh et al., 2018) | 21.18 | 13.45 | 12.03 | 91.18 | 63.88 |
| From sequence | 21.29 (monomer) | 13.99 (monomer) | 15.82 (monomer) | 96.62 (dimer) | 63.49 (monomer) |
| Rigid body modeling |  |  |  |  |  |
| CORAL |  |  |  |  |  |
| Symmetry | - | P1 | P1 | - | - |
| s-range for fit (nm<sup>-1</sup>) | - | 0.419 – 4.603 | 0.321 – 4.09 | - | - |
| χ<sup>2</sup>, CorMap P-value | - | 1.293 / 0.00000002 | 1.768 / 0.000000000000004 | - | - |
| EOM |  |  |  |  |  |
| Symmetry | P1 | - | - | P1 | P1 |
| s-range for fit (nm<sup>-1</sup>) | 0.077 – 4.988 | - | - | 0.140 – 4.340 | 0.099 – 4.895 |
| χ<sup>2</sup>, CorMap P-value | 1.062 / 0.000013 | - | - | 1.179 / 0.160 | 1.335 / 0.0122 |

| SASBDB accession codes (Kikhney et al., 2020) | SASDWV3 | SASDWT3 | SASDWW3 | SASDWS3 | SASDWU3 |
| --- | --- | --- | --- | --- | --- |
| **Software** |  |  |  |  |  |
| ATSAS Software Version (Manalastas-Cantos et al., 2021) | 3.0.5 |  |  |  |  |
| Primary data reduction | PRIMUS (Konarev et al., 2003) |  |  |  |  |
| Data processing | GNOM (Svergun, 1992) |  |  |  |  |
| Rigid body modelling | CORAL (Petoukhov et al., 2012) |  |  |  |  |
| Flexibility ensemble modelling | EOM (Bernadó et al., 2007; Tria et al., 2015) |  |  |  |  |
| Model visualization | PyMOL (PyMOL, 2022) |  |  |  |  |

$s = 4\pi \sin(\theta)/\lambda$ , $2\theta$ – scattering angle, n.d. not determined

**Table S3:** Best matches from DALI searches against PDB25 (cutoff Z value ≥7)

| crTP<sub>part2, RnaH</sub> |  |  |  |  |  |  |  |
| --- | --- | --- | --- | --- | --- | --- | --- |
| No | Chain | Z | rmsd | lali | nres | %id | PDB Description |
| 1 | 5hwi-A | 10.3 | 2.0 | 101 | 229 | 13 | GLUTATHIONE-SPECIFIC GAMMA-GLUTAMYL CYCLOTRANSFERASE |
| 2 | 2qik-A | 9.2 | 2.2 | 94 | 269 | 18 | UPF0131 PROTEIN |
| 3 | 2i5t-A | 9.1 | 2.6 | 100 | 169 | 15 | PROTEIN C7ORF24 |
| 4 | 2jqv-A | 8.4 | 2.4 | 97 | 165 | 11 | AIG2 PROTEIN-LIKE |
| 5 | 3juc-A | 8.1 | 2.5 | 97 | 150 | 14 | AIG2-LIKE DOMAIN-CONTAINING PROTEIN 1 |
| 6 | 1v30-A | 7.8 | 2.3 | 89 | 118 | 15 | HYPOTHETICAL UPF0131 |
| 7 | 6ky1-C | 7.2 | 1.9 | 77 | 152 | 12 | GLUTATHIONE-SPECIFIC GAMMA-GLUTAMYL CYCLOTRANSFERASE |
| crTP<sub>part2, ArgC</sub> |  |  |  |  |  |  |  |
| No | Chain | Z | rmsd | lali | nres | %id | PDB Description |
| 1 | 5hwi-A | 9.3 | 1.7 | 86 | 229 | 17 | GLUTATHIONE-SPECIFIC GAMMA-GLUTAMYL CYCLOTRANSFERASE |
| 2 | 2i5t-A | 8.6 | 1.6 | 83 | 169 | 23 | PROTEIN C7ORF24 |
| 3 | 2qik-A | 8.0 | 1.8 | 79 | 269 | 23 | UPF0131 PROTEIN YKQA |
| 4 | 3juc-A | 7.4 | 2.0 | 81 | 150 | 17 | AIG2-LIKE DOMAIN-CONTAINING PROTEIN 1 |
| 5 | 2jqv-A | 7.4 | 1.7 | 80 | 165 | 21 | AIG2 PROTEIN-LIKE |

**Table S4:** Protein structures available for the γ-glutamyl cyclotransferase-like superfamily.

| Protein | UniProt | PDB code of the structure of the natural protein | PDB code of additional structures | References |
| --- | --- | --- | --- | --- |
| *Arabidopsis thaliana* At5g39720.1 | Q9FIX2 | 2g0q |  | (Lytle et al., 2006) |
| *Arabidopsis thaliana* At3g28950.1 | Q9MBH1 | 2jqv |  | (de la Cruz et al., 2008) |
| *Homo sapiens* γ-glutamylamine cyclotransferase (GGACT) | Q9BVM4 | 3jub | 3juc (Complex w/ 5-oxoproline)<br>3jud (E82Q mutant) | (Oakley et al., 2010) |
| *Mus musculus* aig2-like protein (a2ld1, ggact, mgc7867) | Q923B0 | 1vkb (crystal)<br>2kl2 (NMR) |  | (Klock et al., 2005)<br>(Serrano et al., 2010) |
| *Pyrococcus horikoshii* PH0828 | O58558 | 1v30 |  | (Tajika et al., 2004) |
| *Escherichia coli* YtfP | P0AE48 | 1xhs (NMR) |  | (Aramini et al., 2007) |
| *Bacillus subtilis* YkqA | P39759 | 2qik |  | - |
| *Homo sapiens* | O75223 | 2pn7 | 2rbh (E98A mutant)<br>3cry (E98Q mutant) | (Oakley et al., 2008) |
| γ-glutamyl cyclotransferase (GGCT) | | 2i5t | | 07Bae (Bae et al., 2008) |
| *Saccharomyces cerevisiae* ChaC2 | P32656 | 5hwi (seleno-methionine mutant) | 5hwk (benzoic acid complex) | (Kaur et al., 2017) |
| *Homo sapiens* Glutathione-specific γ-glutamylcyclotransferase 2 (ChaC2) | Q8WUX2 | 6k95 | 6ky0 (E74Q mutant)<br>6ky1 (E83Q mutant) | (Nguyen et al., 2020) |

**Table S5:**
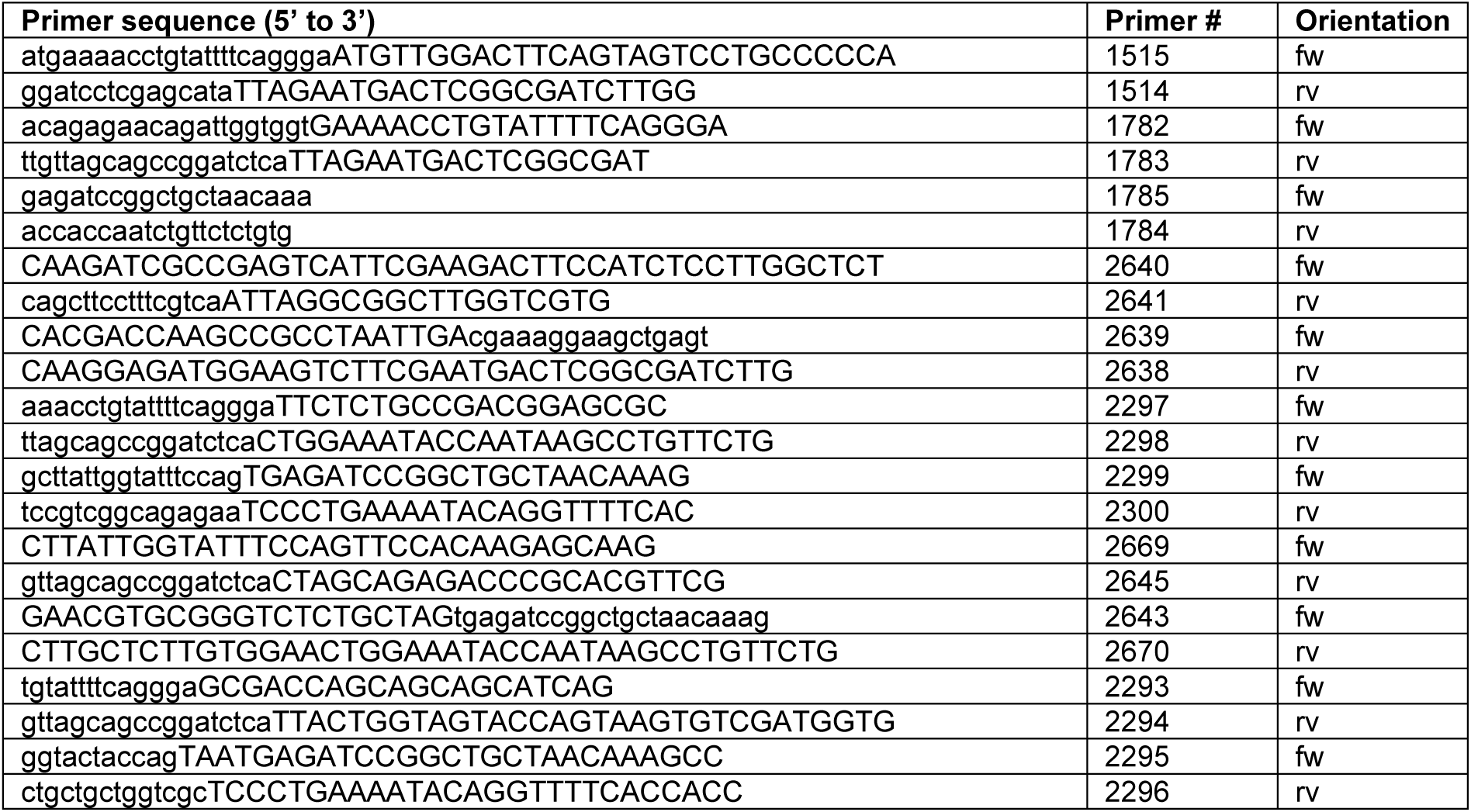
Nucleotide sequences of primers used in this study. The table provides sequence, internal primer number and indicates forward (fw) or reverse (rv) orientation for each primer.

